# Rethinking symbiotic metabolism: trophic strategies in the microbiomes of different sponge species

**DOI:** 10.1101/2021.08.28.458021

**Authors:** I Burgsdorf, S Sizikov, V Squatrito, M Britstein, BM Slaby, C Cerrano, KM Handley, L Steindler

**Author notes:** Corresponding author: Laura Steindler Department of Marine Biology, Leon H. Charney School of Marine Sciences, University of Haifa, 199 Aba Khoushy Ave, Mount Carmel, Haifa, 3498838, Israel, /Fax: /.

## Abstract

In this study we describe the major lithoheterotrophic and autotrophic processes in 21 microbial sponge-associated phyla using novel and existing genomic and transcriptomic datasets. We show that a single gene family, molybdenum-binding subunit of dehydrogenase (*coxL*), likely evolved to benefit both lithoheterotrophic and organoheterotrophic symbionts, through adaptation to different inorganic and organic substrates. We show the main microbial carbon fixation pathways in sponges are restricted to specialized symbiotic lineages within five phyla. We also propose that sponge symbionts, in particular Acidobacteria, are capable of assimilating carbon through anaplerotic processes. However, the presence of symbionts genomically capable of autotrophy does not inform on their actual contribution to light and dark carbon fixation. Using radioisotope assays we identified variability in the relative contributions of chemosynthesis to total carbon fixation in different sponge species. Furthermore, the symbiosis of sponges with two closely related Cyanobacteria results in outcomes that are not predictable by analysis of -*omics* data alone: *Candidatus* Synechococcus spongiarum contributes to the holobiont carbon budget by transfer of photosynthates, while *Candidatus* Synechococcus feldmannii does not. Our results highlight the importance of combining sequencing data with physiology to gain a broader understanding of carbon metabolism within holobionts characterized by highly diverse microbiomes.

## Introduction

Microbes can autotrophically assimilate inorganic carbon (Ci) via seven pathways including the Calvin-Benson-Bassham (CBB) cycle, the reductive tricarboxylic acid (rTCA) cycle, the Wood-Ljungdahl pathway (WL), and the 3-hydroxypropionate/4-hydroxybutyrate (3-HP/4-HB) cycle [1]. Additionally, various solo-acting enzymes can be involved in Ci assimilation without being part of *sensu stricto* carbon fixation. For instance, Ci assimilation in anaplerotic reactions was proposed to be abundant among marine planktonic heterotrophs [2–4]. Anaplerotic reactions undertaken by pyruvate (PYC) and phosphoenolpyruvate (PPC) carboxylases often occur at low levels to replace intermediates of the tricarboxylic acid (TCA) cycle. However, enhanced anaplerotic Ci assimilation was reported in marine planktonic lithoheterotrophs that combine organotrophy with the additional use of inorganic electron donors [4]. Malic enzyme (MEZ) was also shown to operate in the carboxylating (anaplerotic) direction, playing an essential role in the intracellular survival of the pathogen *Mycobacterium tuberculosis* [5, 6], in planktonic stages of *Pseudomonas aeruginosa* PAO1 [7] and in deep-sea Alphaproteobacteria [8].

Sponges (phylum Porifera) are ancient cosmopolitan filter-feeders [9, 10]. They play an important role in nutrient recycling by transforming Dissolved Organic Matter (DOM) into detrital Particulate Organic Matter, thereby making it available for other invertebrates in nutrient poor environments [11, 12]. Sponge-associated microbial communities include more than 60 bacterial and archaeal phyla [13], which are specific to their hosts [14–17]. These sponge-associated symbionts can be categorized based on their nutrition strategies, for instance (photo- and chemo-) autotrophic, organoheterotrophic and lithoheterotrophic. Heterotrophic symbionts can contribute up to 87% of the total sponge holobiont DOM assimilation [18], while autotrophic photosymbionts can contribute to host growth when exposed to light [19]. Beyond photosymbionts, various bacterial and archaeal phyla in sponges harbor mechanisms associated with autotrophic metabolism, *e.g.*, the 3-HP/4-HB pathway in Thaumarchaeota, and the rTCA in Nitrospirota, Alphaproteobacteria and Oligoflexia [20–25]. However, carbon assimilation capacities within the sponge microbiome remain under-described, and the contribution of chemoautotrophy to the pool of microbially-fixed carbon in sponges has not yet been tested.

Net primary productivity and stable isotope analyses of the microbial and host sponge fractions showed that different species of symbiotic filamentous and unicellular Cyanobacteria differ in their ability to assimilate and transfer carbon to the host [26]. The unicellular *Parasynechococcus*-like cyanobacterial species are the most commonly reported in sponges [27, 28]. These include *Candidatus* Synechococcus spongiarum, enriched in 28 sponge species around the globe (including *Theonella swinhoei* from this study) [29], and *Candidatus* Synechococcus feldmannii, the symbiont of *Petrosia ficiformis* [30, 31]. The latter symbiosis is facultative, with *P. ficiformis* growing in light environments with *Ca.* S. feldmannii, and in dark-(cave)-environments without it. The heterotrophic microbial community of *P. ficiformis* is functionally and compositionally independent from the presence of *Ca*. S. feldmannii, being nearly identical in both structure and gene expression in specimens with and without this photosymbiont [31, 32]. This suggests that photosynthetically derived carbon may not be the main carbon source for heterotrophic *P. ficiformis*-associated symbionts. In this study we tested whether microbially fixed carbon or rather accumulation of DOM by the host represent the main supply of organic carbon to *P. ficiformis*.

Oxidation of diverse inorganic compounds, such as ammonia, nitrite, sulfide and thiosulfate, can serve as energy source for both autotrophic and lithoheterotrophic microorganisms, and these can also be found within the sponge microbiome [33–36]. Here, we characterized the dominant carbon fixation processes, and identified the energetic sources used by lithoheterotrophs across different microbial species within sponge symbiotic communities. This was achieved through a genomic analysis of 402 symbiotic metagenome-assembled genomes (MAGs) from 10 different sponge species, and 39 metatranscriptomes from the sponge *P. ficiformis*. Further, using radioisotopes, we investigated the contribution of light and dark microbial carbon fixation in two sponge systems (*P. ficiformis* and *T. swinhoei*).

## Materials and methods

### Sponge sampling and microbial DNA purification

Three *P. ficiformis* specimens, 277c, 287ce, and 288c (c, cortex; e, endosome), were collected by SCUBA diving on January 6^th^, 2014, at depths 27, 23 and 15 meters, respectively, at Achziv nature marine reserve, Mediterranean Sea, Israel. Sponge samples were collected in compliance with permits from the Israel Nature and National Park Protection Authority. Microbial DNA was obtained as previously described [37]. Information about collection of additional nine sponge species can be found in Table S1.

### Shotgun sequencing, assembly and binning

Preparation of metagenomic shotgun sequencing KAPA Hyper DNA libraries, sequencing, read trimming and *de novo* assemblies for the three *P. ficiformis* specimens were performed as previously described [37]. 50 genomes were binned using manual methods including visualization of differential coverage information derived from three *P. ficiformis* specimens (see File S1 at https://figshare.com/s/e305160ebd82d21bf151). DNA extraction, and assembly of MAGs from *T. swinhoei* (SP3), *Ircinia variabilis* (142) and *Aplysina aerophoba* (15L) are described in [38] and [21], respectively. Taxonomic affiliation of assembled scaffolds, binning of final MAGs, and relative abundance calculations are described in File S2.

### MAG annotation and completeness estimation

Open Reading Frames were identified using Prodigal v2.6.3 with the metagenome options [39]. Protein sequences were queried against the Clusters of Orthologous Groups (COGs) database (version 2014) as previously described (see File S3 at https://figshare.com/s/786acca3672a820da570) [37]. The amino acid sequences were also searched against the KEGG orthology (KO) database using standalone KofamKOALA 1.3.0 (see File S4 at https://figshare.com/s/a0ff9b495588b99e7c75) [40]. Selected enzymes were annotated using previously published Hidden Markov models (HMM) [41] with individual score thresholds (Table S2) using hmmsearch [42]. Phylogenomic tree construction and taxonomic annotation was done using PhyloPhlAn2 [43] (https://bitbucket.org/nsegata/phylophlan/wiki/phylophlan2), RAxML [44] as previously described [45] and GTDB-Tk v1.3 with release r95 [46]. Trees were visualized using iTol [47]. Completeness and contamination rates of all final MAGs were estimated with checkM version 1.0.7 [48] using lineage_wf.

### Annotation of transcriptomic data

Metatranscriptomes were previously obtained from 39 *P. ficiformis* specimens sampled in Ligurian Sea, Italy (as described in [31]), and quantified by Salmon software [49]. Translated sequences of the assembled and filtered bacterial metatranscriptomes were assigned to the proteins of MAGs assembled from Israeli *P. ficiformis* specimen 277c using blastp 2.2.30+ (*E*-value threshold = 1E−10, identity = 55%). Taxonomic annotation of transcripts was determined based on the best hits (highest bit score and lowest *E*-value). Transcripts with the same function and MAG affiliation were merged prior to analyses. Expression of certain functions in a specific organism (MAG) was additionally confirmed using mapping of the genes against metatranscriptome reads with bbmap tool v 37.62 [50] (minimal identity=0.70, kmer size=13) from the BBtools package (https://jgi.doe.gov/data-and-tools/bbtools/), ( ≥ 5 reads as a threshold). Functional annotation of the metatranscriptomes of *P. ficiformis* against the COG database was done as described previously [37] and search against the KEGG database was done via the GhostKOALA website using the genus_prokaryotes database (August 2020) [51]. Data was analyzed and visualized using the R packages dplyr, tidyr [http://tidyr.tidyverse.org], ggplot2 [52], plotly [53], reshape2 [54] and superheat [55]. A schematic representation of the bioinformatic analyses is available in Figure S1.

### Carbon fixation measurements in *Theonella swinhoei* and *Petrosia ficiformis* with H^14^CO_3_^-^

Photosynthetic (light) and chemosynthetic (dark) fixed carbon was measured in *P. ficiformis* (n=3) and *T. swinhoei* (n=4) specimens. For *P. ficiformis*, experiments were done on sponge cores, for *T. swinhoei*, on whole sponges and sponge cores. In *P. ficiformis* there are sponge areas facing light, that harbor Cyanobacteria (*Ca*. S. feldmannii), and others facing the shade, without Cyanobacteria. We thus used sponge cores from both areas with and without *Ca.* S. feldmannii in our experiments. Based on a pre-experiment on *T. swinhoei* cores, we determined the optimal incubation time of 2 hours (data not shown), used for all follow-up experiments. Next, we incubated cores (*P. ficiformis*) and whole sponges (*T. swinhoei*) in beakers with autoclaved seawater and 1 µl of NaH^14^CO_3_ (ARC, 150922, 1mCi/1ml) tracer for each 10 ml medium, stirring manually every 10 min. Beakers were exposed to light (100-150 and 50 µmol photons m^-2^ s^-1^, for *P. ficiformis and T. swinhoei*, respectively) or to darkness. NaH^14^CO_3_ in the medium was measured every 30 min, by sampling 0.1 ml of seawater from each beaker and transferring to a scintillation vial containing 3 ml of scintillation fluid (UltimaGold®, Perkin-Elmer for *P. ficiformis* and Opti-fluor®, high flash-point LSC cocktail, Packard Bioscience for *T. swinhoei*). At the end of the incubation, each core section was left for 3 minutes on a paper towel to lose excess water, weighted and then transferred to a vial containing 0.5 ml N,N-Dimethylformamide (Sigma), to release labeled fixed carbon from the tissue to the liquid, and 45 µl HCl 20% (Sigma), to release labeled/unlabeled non-fixed carbon. Sponge tissue was disintegrated manually with a plastic homogenizer and the vials were left uncovered for 48 hours in the chemical hood. After this time, 0.1 ml liquid was transferred to a new vial containing 3 ml scintillation fluid. All vials were measured in a liquid scintillation counter. Tri-Carb® 2810TR, Perkin Elmer and Tri-Carb® 1500, Lumitron, Packard Bioscience were used to measure the Discharges Per Min (DPM) for *P. ficiformis* and *T. swinhoei*, respectively. A detailed explanation about sponge sampling and core preparations for C fixation experiments is available in the File S2.

## Results and Discussion

Overall, 47 bacterial and 3 archaeal MAGs belonging to 14 phyla were recovered from three *P. ficiformis* specimens, and are estimated to be 62 to 100% complete, with 0 to 5.5% contamination (Table S3). These genomes represent 38-41% of the assembled data and were investigated together with 8 additionally assembled MAGs from *T. swinhoei* (SP3), *Ircinia variabilis* (142) and *A. aerophoba* (15L) (Table S3), and additional 344 MAGs from previous studies [21, 24, 61–66, 25, 29, 37, 56–60] (Table S1) to identify all the dominant autotrophic and lithoheterotrophic processes found in sponge symbionts.

### Autotrophy in sponge symbionts

We investigated the presence of known prokaryotic carbon fixation mechanisms among 402 sponge-associated MAGs derived from 10 sponge species (Figure 1, Figure S2, Tables S1, S3 and S4). The metabolic capacities of the sponge-associated symbionts are presented in Table S4 and their predicted trophic lifestyles, are summarized in Figure 2. Accordingly, we found that the autotrophic pathways 3-HP/4-HB, CBB and rTCA were mainly restricted to Cyanobacteria, Tectomicrobia, Nitrospirota, and Thaumarchaeota phyla and two gammaproteobacterial orders.

**Figure 1.**
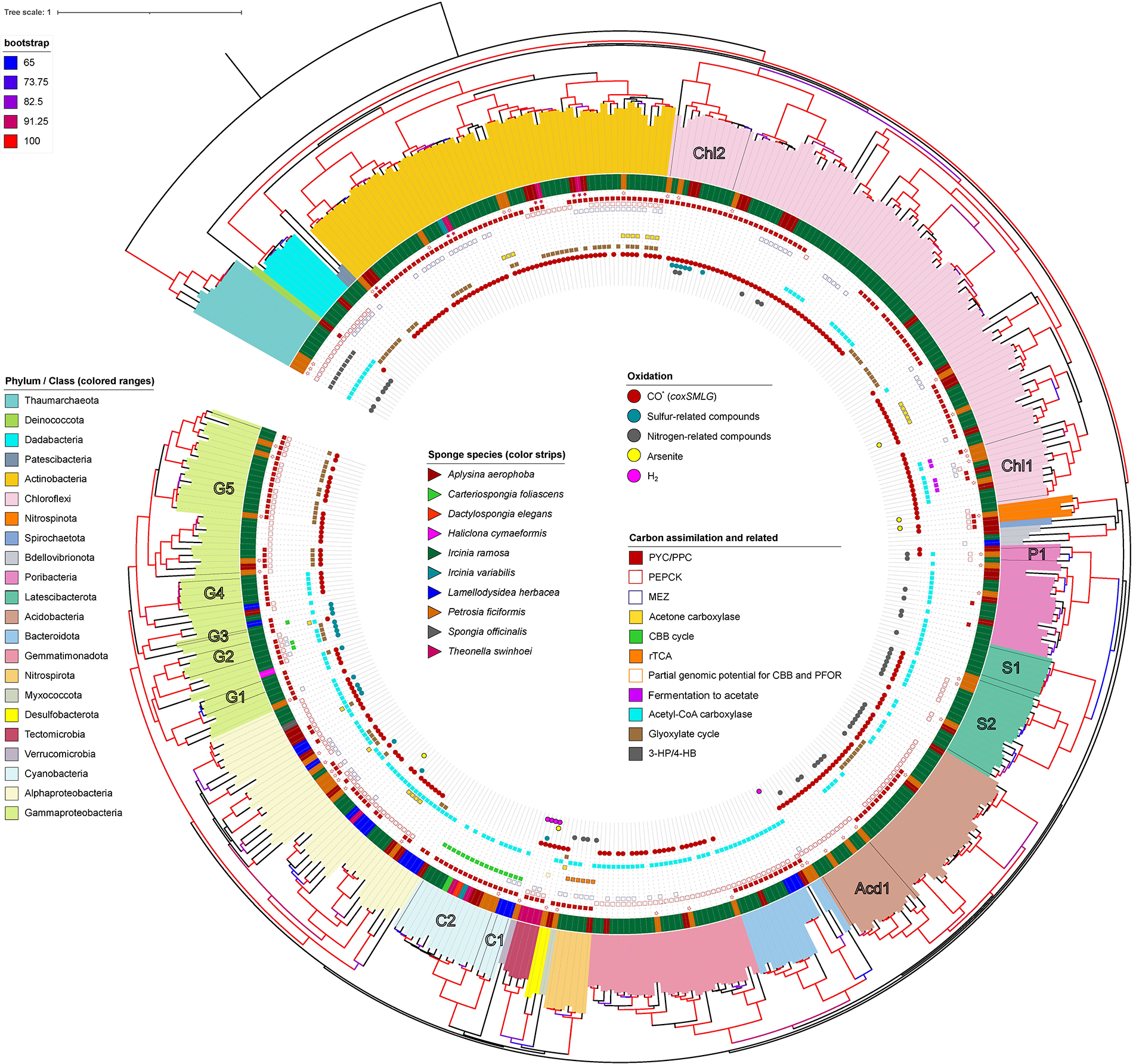
Phylogenomic tree showing the distribution and diversity of carbon assimilation and energy production pathways across microbial symbiont taxonomy and host species. The phylogenomic tree (N=399 MAGs) was constructed based on concatenated universal markers (PhyloPhlAn2). Labels marked with a hollow star are MAGs assembled in this study from the *P. ficiformis* specimen 277c. Labels marked with a colored star are eight MAGs assembled from the *A. aerophoba* specimen 15, *T. swinhoei* specimen SP3 and *I. variabilis* specimen 142. The tree is rooted to the Archaea group. Figure S6 represents an enhanced version (MAGs names are displayed) of this tree. Acd1, class Vicinamibacteria, order *Vicinamibacterales*, family UBA8438. C1, order *Cyanobacteriales*, family *Desertifilaceae*. C2, order *Synechococcales*, family *Cyanobiaceae*. CHL1, class Dehalococcoidia, order UBA3495. CHL2, class Anaerolineae, order SBR1031. G1, order GCA-2729495. G2, order UBA10353, family LS-SOB. G3 (single MAG), order UBA4575. G4, order *Pseudomonadales, Pseudohongiellaceae* family. G5, order *Pseudomonadales,* HTCC2089 family. P1, class and order WGA-4E, unknown family. S1, unknown class. S2, UBA2968 class and order. Poribacteria_ADFK02.1_Kamke_2014, Poribacteria_AQPC01.1_Kamke_2014 and Poribacteria_ASZM01.1_Kamke_2014 were excluded from the phylogenomic tree due to incomplete marker genes set. *CO is not always a target molecule for the *coxSMLG* complex as it was shown here for *Poribacteria*.

**Figure 2.**
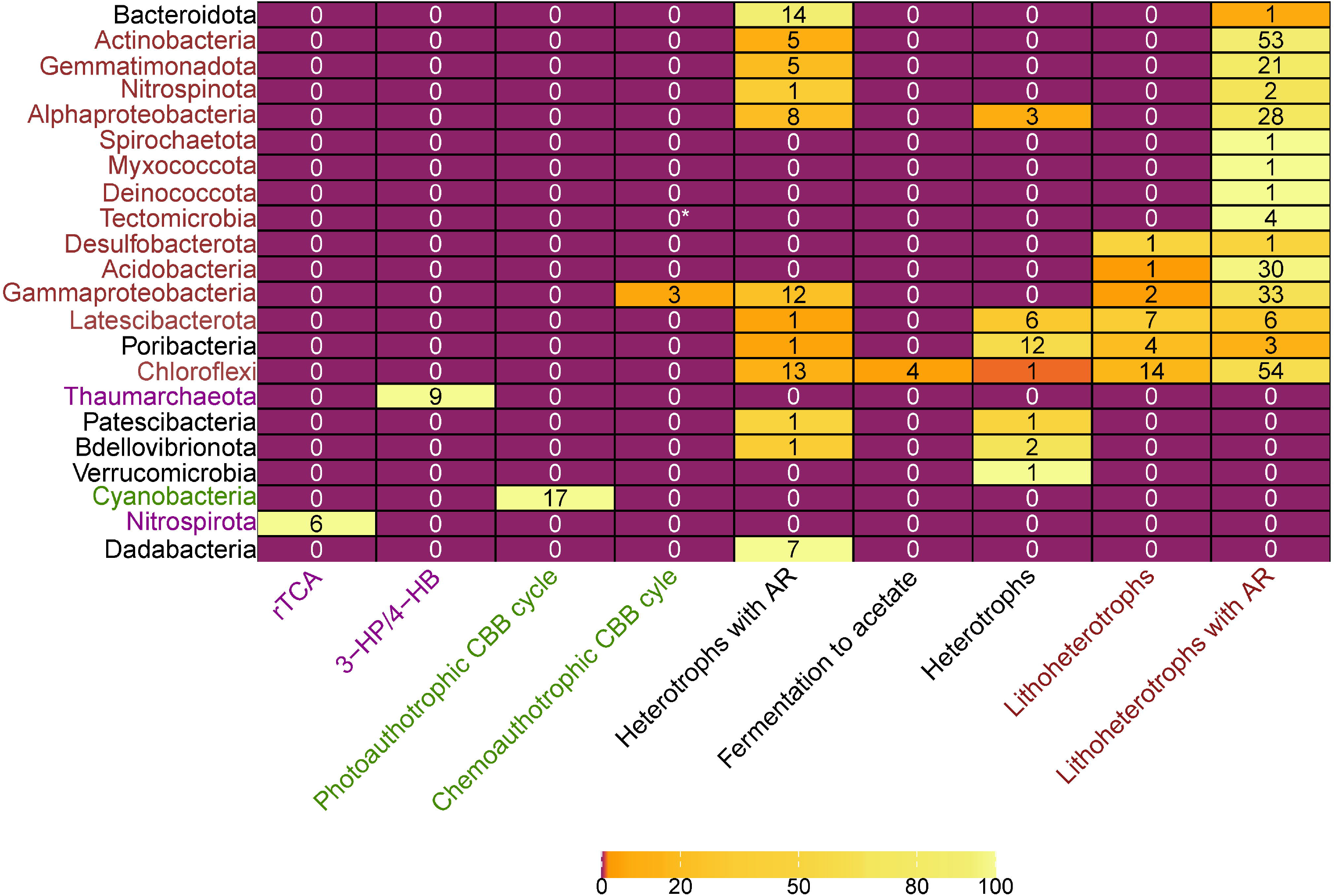
Predicted lifestyle for different taxonomic groups (Phylum/Class) of sponge symbionts. Heat map represents percentage of genomes with predicted lifestyle, text represents number of MAGs. The colors of the MAGs correspond to the most abundant lifestyle: lithoheterotrophs (red), autotrophs implementing CBB (green) and other chemoautotrophs (violet). The relevant functions can be found in Table S4. Here, heterotroph means organoheterotroph. AR, anaplerotic reaction. *MAGs of Tectomicrobia (Entotheonellia) class possess incomplete genomic potential for utilization of CBB pathways.

RuBisCO-related genes were identified in all 16 cyanobacterial and 3 (out of 47) gammaproteobacterial genomes. Gammaproteobacterial MAGs from *P. ficiformis* lacked RuBisCO-related genes, or evidence of any other C-fixation pathways, and thus likely pursued a heterotrophic lifestyle. Yet, within these sponge species, 2 out of 5 gammaproteobacterial MAGs exhibited genomic potential for CO oxidation via carbon monoxide dehydrogenase (CODH), which can serve as energy source [67]. In *I. ramosa,* Gammaproteobacteria clades G2 (order UBA10353 and family LS-SOB) and G3 (order and family UBA4575) had genes for thiosulfate oxidation (Table S4), which can fuel carbon fixation (Figure 1, Figure S2) through RuBisCO (also found in three MAGs within G2 and G3 clades, File S2), indicating the potential for chemoautotrophy. Gammaproteobacteria G1, G4 and G5 did not have this pathway, yet encoded for CODH (*coxSML* and *coxG*) (Figure 1, Figure S2). Taken together, data show that gammaproteobacterial symbionts include two trophic groups: chemoautotrophs and lithoheterotrophs.

The filamentous Entotheonellia (phylum Tectomicrobia) found in the sponge *T. swinhoei* [61, 68, 69], were identified as chemoautotrophs based on the presence of a large cohort of CBB related genes [61]. We did not detect RuBisCO in the Tectomicrobia genomes (Table S4), which may be due to MAG incompleteness. Energy for carbon fixation in this phylum may be provided by oxidation of multiple inorganic donors, providing metabolic versatility to shifting environmental conditions within the host [70, 71]. Inorganic donors (and mechanisms for oxidation) include CO (CODH), H_2_ (3b group hydrogenase), thiosulfate (Sox complex), and possibly even arsenite (AoxAB) (Table S4). Presence of arsenic was reported in sponges, at highest concentrations in *T. swinhoei* [72]. Oxidation of arsenite may have a dual function: energy source [73, 74], as well as detoxification of the highly toxic arsenite to arsenate [75]. Calcium arsenate was in fact observed inside intracellular structures of filamentous Tectomicrobia [68]. Arsenite oxidation is not necessarily limited to filamentous Tectomicrobia, in fact we also found the arsenite oxidase genes (*aoxAB*) in Alphaproteobacteria, Chloroflexi, and Nitrospinota MAGs (Figure 1, Table S4).

The pyruvate synthase or pyruvate:ferredoxin oxidoreductase (PFOR, EC 1.2.7.1), which is required for the rTCA cycle, can also serve different non-autotrophic functions, such as energy production through fermentation of pyruvate to acetate. For example, sediment Chloroflexi harboring PFOR and acetyl-CoA synthetase [EC 6.2.1.1]) were predicted to biosynthesize ATP using this pathway [76]. Here, PFOR was identified in five Chloroflexi MAGs (from *A. aerophoba*, *P. ficiformis* and *I. ramosa*) that lack carbon fixation pathways and may serve an energy production role. Accordingly, a high abundance of acetyl-CoA synthetase (COG1042) was previously detected in diverse sponge microbial metagenomes [77]. We therefore hypothesize that in the studied sponges ATP production involving pyruvate conversion to acetyl-CoA (by PFOR), coupled with acetate formation (by acetyl-CoA synthetase), occurs in five specialized, sponge-associated Chloroflexi (Figure 1, Figure S2, Table S4).

### Lithoheterotrophy and metabolism of inorganic compounds in sponge symbionts

We and others have detected genes for oxidation of diverse inorganic compounds, such as CO, nitrite, ammonia, and thiosulfate in the sponge microbial community [33–36]. Further, we reveal the potential for hydrogen oxidation, specifically by the Tectomicrobia derived from *T. swinhoei* and in a Bacteroidetes MAG from *P. ficiformis* (Figure 1, Table S4, Figure S2). Nitrogen processing by diverse members of the sponge microbiome has been analyzed in several studies (*e.g.*, [33, 36]). It was suggested that ammonia oxidation in sponges is uniquely performed by Thaumarchaeota [36]. Nitrite can be oxidized to nitrate by members of Nitrospirota, Alphaproteobacteria and Gammaproteobacteria symbionts [35]. We speculate that oxidation of nitrite to nitrate may only be carried out by Nitrospirota, rather than also by Proteobacteria, as previously proposed [35]. We base this speculation on a stricter annotation of the genes involved (*nrxAB*) using both HMM profiles and KEGG annotations (File S2).

Orthologues of *amoABC/pmoABC* genes (involved in ammonia and/or methane oxidation [78, 79], Table S2) were here found also in Desulfobacterota (unclassified Deltaproteobacteria based on NCBI taxonomy), specifically in two MAGs deriving from *A. aerophoba* and *P. ficiformis* (Table S4, File S2). Based on sequence similarity, we predict that *amoABC/pmoABC* of Desulfobacterota are involved in methane oxidation (File S2). Besides methane to methanol oxidation (EC 1.14.18.3), these MAGs also have the potential to further oxidize methanol to formaldehyde (EC:1.1.2.10) (Table S4). Genomic potential for methane to formaldehyde oxidation was previously discovered in sponges, but was not affiliated with members of Desulfobacterota [36, 80]. *amoABC/pmoABC* subunits were shown to be also expressed within the sponge *P. ficiformis* (details provided below).

CO-oxidizing bacteria are lithoheterotrophs common in sponge microbiomes. Large (CoxL, COG1529) and middle (CoxM, COG1319) subunits of the molybdenum-rich aerobic form of CODH (Mo-CODH) are highly overrepresented in sponge-associated *versus* seawater microbial metagenomes [34]. Mo-CODH has been identified in gamma and alphaproteobacterial sponge symbionts [33, 34] and found to be expressed among phylogenetically diverse symbionts including Actinobacteria, Chloroflexi and Proteobacteria [81]. Yet the function of CoxL is variable, and its homologues are not solely responsible for CO oxidation. In fact, CoxL was shown to comprise two different forms (I and II), with form II (putative *coxL*) being involved in functions alternative to CO oxidation [67]. To establish the extent to which CO oxidation is abundant in sponge symbionts, determine potential alternative substrates beyond CO, and provide this information at a taxonomic level, we set the following criteria: (i) genomic potential for CO oxidation was based on the presence of 4 subunits (*coxSMLG*) within MAGs, (ii) substrate specificity was based on clustering and reannotation of 2,406 translated *coxL* genes against the KO database and on reannotation of transcripts, (iii) taxonomy of transcripts was defined according to MAG-affiliation.

The Mo-CODH complex was found in 64% of all analyzed symbionts (Figure 1, Table S4), suggesting that CO oxidation is the most abundant process related to a lithoheterotrophic lifestyle in sponge symbionts. Overall, more than half of the protein sequences annotated as CoxL COG1529 belonged to Actinobacteria (29%) and Chloroflexi (22%), while Tectomicrobia and Actinobacteria had the highest average number of *coxL* genes (associated COG1529) per genome (Average = 29, SD = 5 and Average = 12, SD = 4, respectively) (Figure 3A). Among the two largest clusters, one is predicted to function as CO dehydrogenase (the mostly-orange cluster dominated by Actinobacteria, Figure 4), while the second large cluster could not be linked to any known function (the predominantly black cluster, where Chloroflexi prevail). Additional substrates for CoxL are likely isoquinaline (mostly violet cluster, dominated by Gammaproteobacteria) and nicotinate (green cluster, where Gemmatimonadetes and Chloroflexi prevail). Results therefore suggest that sponge symbionts can gain electrons from CO (lithoheterothrophs) and organic molecules (*e.g.*, isoquinaline and nicotinate; organoheterotrophs) via genes related to a large orthologous group – CoxL COG1529. Nevertheless, the substrate for more than half of the proteins annotated as CoxL COG1529 in sponge symbionts, remains unknown (black dots, Figure 4; N/A in Figure 3B).

**Figure 3.**
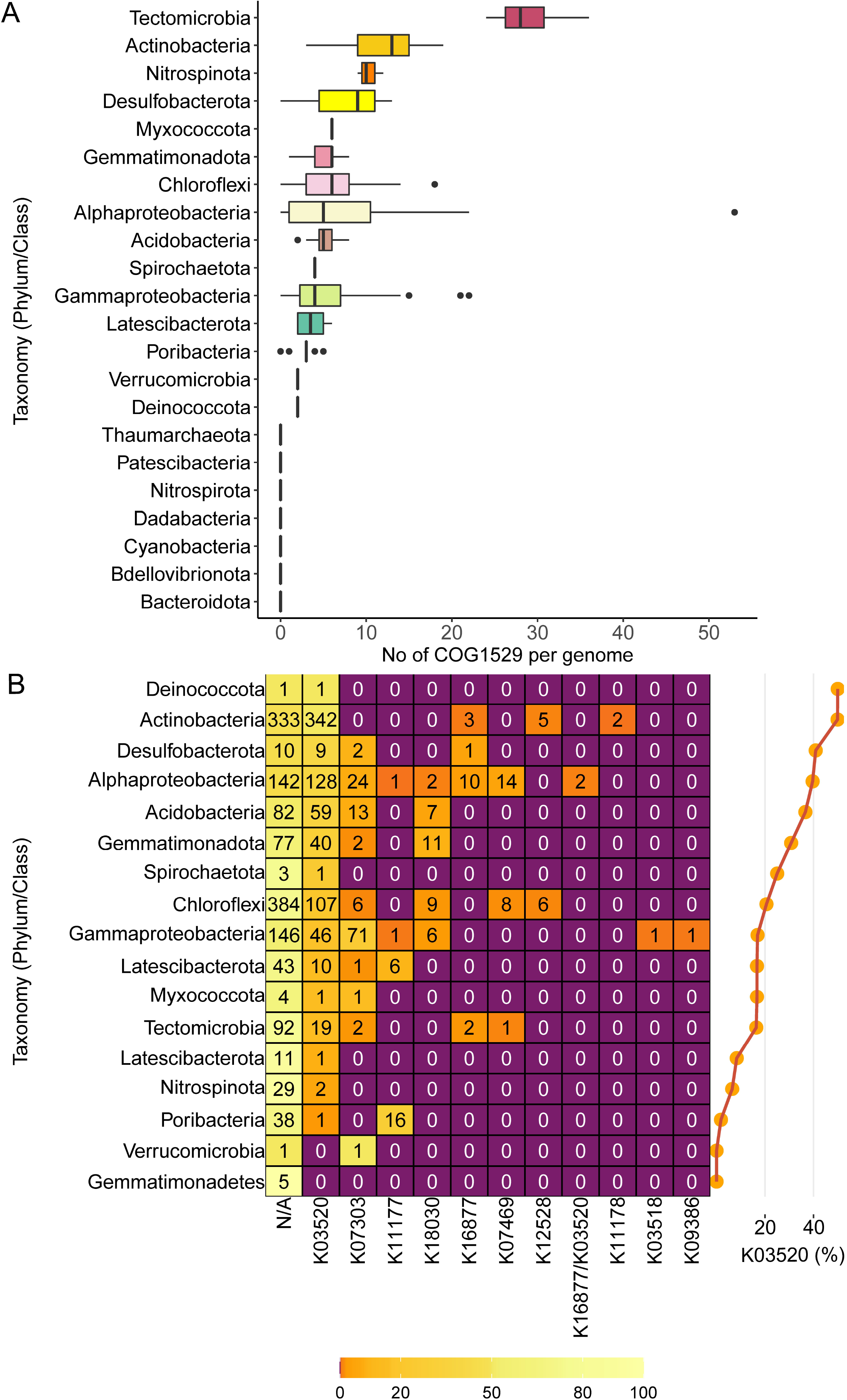
Functional diversity and distribution of COG1529 orthologs across symbiotic bacterial phyla. (A) Number of proteins annotated as COG1529 per genome in different taxonomic groups (Phylum/Class) of sponge symbionts. (B) Functional diversity of the COG1529 orthologous group. Heat map represents percentages of genes with various functions (KEGG annotation) for different taxonomic groups (Phylum/Class) of sponge symbionts. Text represents number of sequences. Percentages of CO-oxidizing *coxL* (K03520) out of total COG1529 are presented on the right. K03520, CO; K07303, isoquinoline; K11177, xanthine; K18030, nicotinate; K16877, 2-furoyl-CoA; K07469, aldehyde; K12528, selenate; K11178, xanthine; K03518, CO; K09386, CO (KO annotation, target molecule).

**Figure 4.**
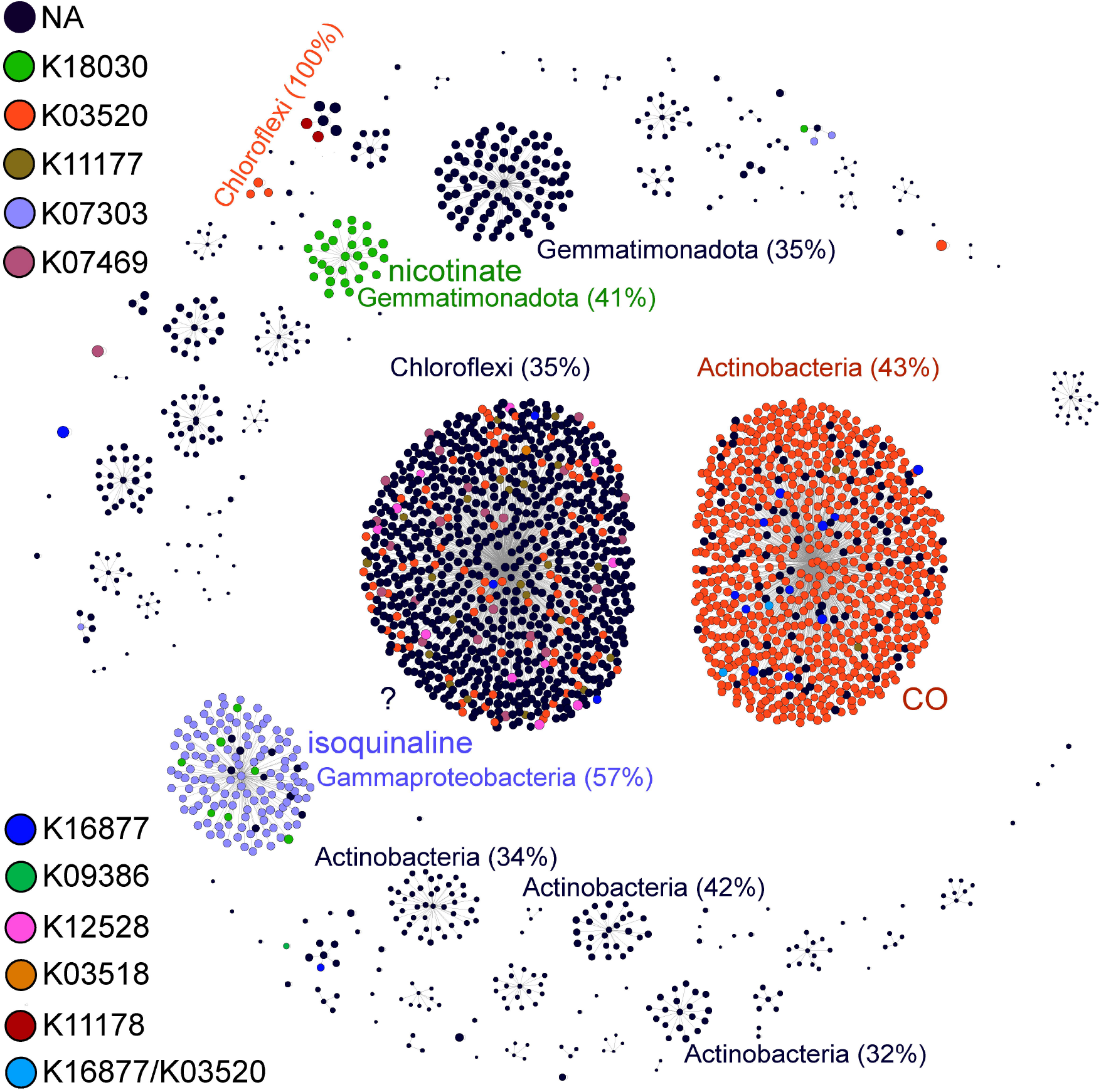
Taxonomic affiliation and hypothesized substrate for CoxL (COG1529) across diversity of sponge-associated MAGs (N=402). Visualization of sequence-based clustering of 2406 proteins annotated as COG1529. Size of the dots is proportional to the length of the protein (in the range of 35-1250 amino acids, average = 682, SD = 207 amino acids). 720 out of 867 sequences forming the largest group (the predominantly black cluster) have unknown function. 674 out of 784 sequences forming the second largest group (the predominantly orange cluster) were annotated as Mo-binding subunit of the CO dehydrogenase (K03520).

While we showed here an extensive incidence of CODH within the sponge microbiome, some phyla were found to lack this functional capacity. Specifically, phyla with inherently autotrophic lifestyles (Cyanobacteria, Nitrospirota, and archaeal Thaumarchaeota) (Figure 2) and phyla specialized in the degradation of polysaccharide residues (Bacteroidota [82, 83], Dadabacteria [84], and Verrucomicrobia [45, 85]), which did not contain CODH (Figure 3B). An exception are the 2 out of 17 Poribacteria, also characterized as degraders of diverse carbohydrate sources originating in the sponge matrix [36, 59, 86], which harbored CODH (Clade P1 in Figure 1). However, CoxL in these two Poribacteria might function in the oxidation of xanthin (see below), and they may thus not have a lithoheterotrophic lifestyle. Mo-CODH should be distinguished from Nickel-CODH, which relates to the anaerobic WL pathway. The latter was previously reported in sponge symbionts [21–23, 25], but based on our analysis (combining KEGG, COG and HMM profiles annotations, Table S2), we conclude that Nickel-CODH, and thus the WL pathway, is absent in the sponge microbiome.

Taken together results indicate that the presence of CoxL COG1529 in sponge symbionts can relate to both CO oxidation (as a part of CODH complex), and thus to a lithoheterotrophic lifestyle, or the oxidation of different organic substrates. Similarly to other symbiotic systems, including the human gut and legumes [67, 87], CO-oxidizing bacteria appear to have an essential role in the sponge holobiont. The potential sources for CO in sponges may include photoproduced CO derived from the ambient seawater [88, 89] and biological hemoprotein degradation via heme oxygenase (HO) activity [67, 87, 90]. Genomic potential for hemoprotein synthesis, transport and oxidation here found in specific members of the sponge microbiome, and suggested as a potential CO source in sponges, is discussed in File S2.

### Gene expression of carbon fixation and energy production pathways: case study of *P. ficiformis*

To study the activity of key processes related to carbon fixation and energy production from oxidation of inorganic molecules, we conducted a genome-informed metatranscriptomic analysis of the *P. ficiformis*-associated community. We linked 50 MAGs (Table S3) with an assembled metatranscriptome dataset derived from 39 *P. ficiformis* specimens. 35 % of transcripts aligned to protein sequences.

Our gene expression results confirm the results derived from the wider MAG analysis described above on the importance of CO oxidation in sponge symbionts, and further corroborate that specific sub-orthologs of COG1529 might provide symbionts with the ability to utilize alternative organic electron donors. The latter may be part of the DOC (or its residues) that is concentrated by the host’s filtration activity [91]. Similar to other sponge species, results show CO-oxidizing bacteria were highly abundant in the sponge *P. ficiformis,* with more than half of the MAGs (64%, n=50) harboring CODH (Figure 1, Table S4), and with all MAGs affiliated to Actinobacteria (n=13), Acidobacteria (n=4), and Chloroflexi (n=9) harboring CODH. Here, we tested how the widespread genomic potential for CO oxidation relates to its expression across different symbiotic microbial phyla.

We confirmed the expression of CO dehydrogenase (K03520) among eight phyla including Acidobacteria, Actinobacteria, and Chloroflexi (Figure 5A). Acidobacteria and Chloroflexi expressed all four subunits of CODH, while Actinobacteria, Desulfobacterota, and Latescibacterota did not express the *coxG* subunit. The *coxG* gene was also absent from the form 1 (*bona fide* CO dehydrogenase) Mo-CODH from the chemoautotroph *Alkalilimnicola ehrlichei* MLHE-1 [67], suggesting that the presence of this gene is not crucial for CO oxidation [92, 93]. Interestingly, while the auxiliary subunits of CODH, affiliated to poribacterial MAGs, were expressed, the large CO-oxidizing subunit was absent in the representatives of this phylum. This may be explained by the functional annotation of the CODH complex as xanthine dehydrogenase (EC 1.17.1.4) in Poribacteria (Figures 3B, 5A and 6), suggesting that this phylum does not oxidize CO in sponges. Functional and taxonomic specialization for certain subgroups of COG1529 was also observed for additional members of the *P. ficiformis* symbiotic microbial community (Figure 5). For instance, a suborthologous group annotated as a subunit of xanthine dehydrogenase (K13482) was exclusively linked to a single actinobacterial MAG (Actino_4, class *Acidimicrobiia*, order UBA5794, family SZUA-232), and a nicotinate dehydrogenase subunit (K18030) was linked to Gammaproteo_5 (order *Pseudomonadales*, family HTCC2089) (Figure 6).

**Figure 5.**
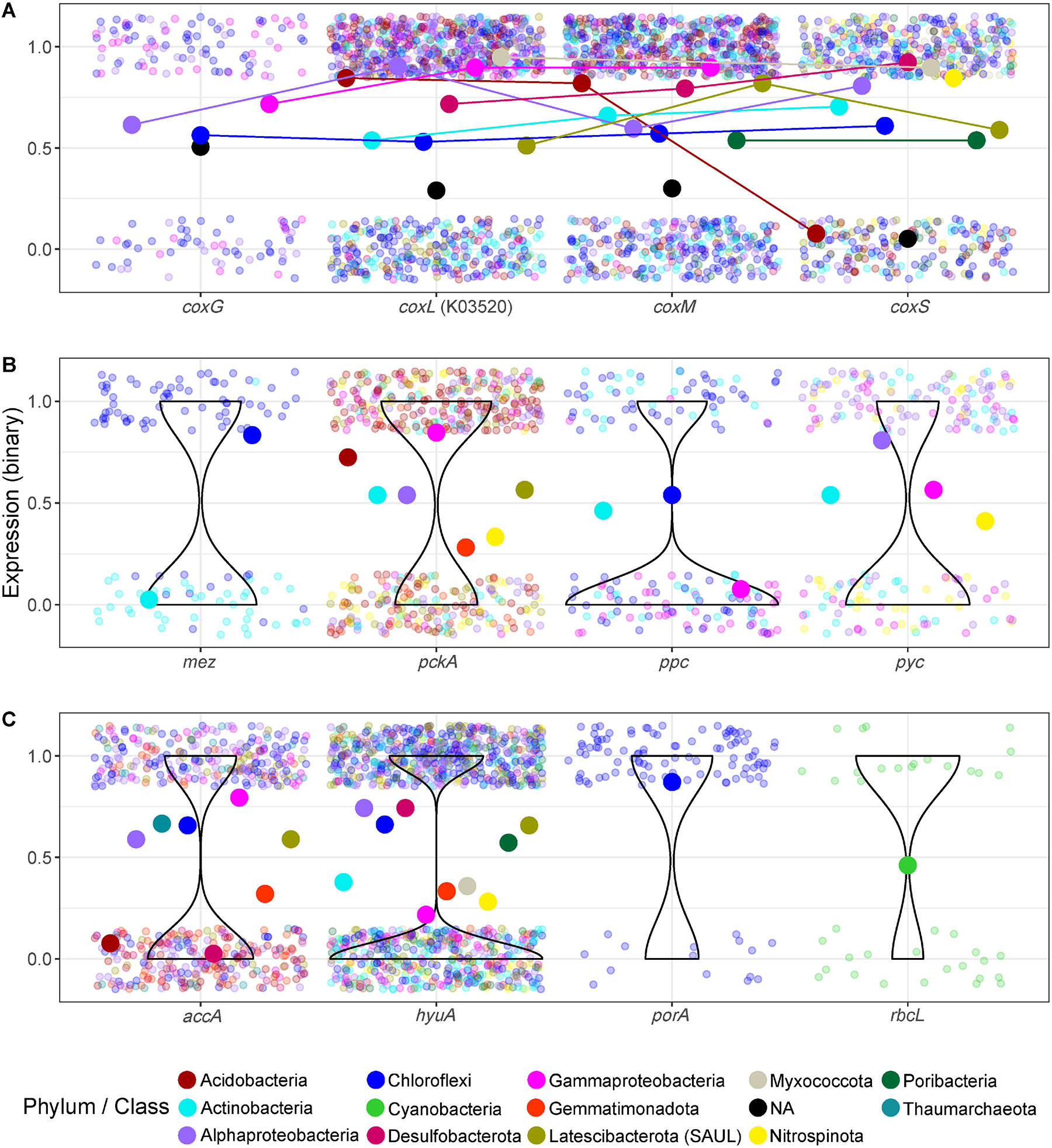
Expression of carbon assimilation and CO oxidation-related functions in the different phyla of *P. ficiformis* symbionts. The analyses are based on cumulative binary (1 – expressed, 0 – not expressed) expression of transcripts (N=39 transcriptomes). Transcripts with the same function and MAG affiliations are merged. (A) the four subunits of CODH (subunits with the same taxonomy are connected by lines), (B) anaplerotic fixation, and (C) carbon assimilation genes. Taxonomy of transcripts was assigned if the transcript was linked to the gene of the assembled MAG. Larger dots represent proportion of expression across samples for a certain taxonomy group (Phylum/Class). Transcripts with not assigned (NA) taxonomy (not linked to any assembled MAG) are given as black dots representing mean values (A) or violin plots representing overall distribution of transcripts (B and C). Genes: *mez*, malic enzyme; *pckA*, phosphoenolpyruvate carboxykinase; *ppc*, phosphoenolpyruvate carboxylase; *pyc*, pyruvate carboxylasev *accA*, subunit of acetyl-CoA carboxylase; *hyuA*, subunit of acetone carboxylase; *porA*, subunit of pyruvate synthase (PFOR); *rbcL*, large subunit of RuBisCO.

**Figure 6.**
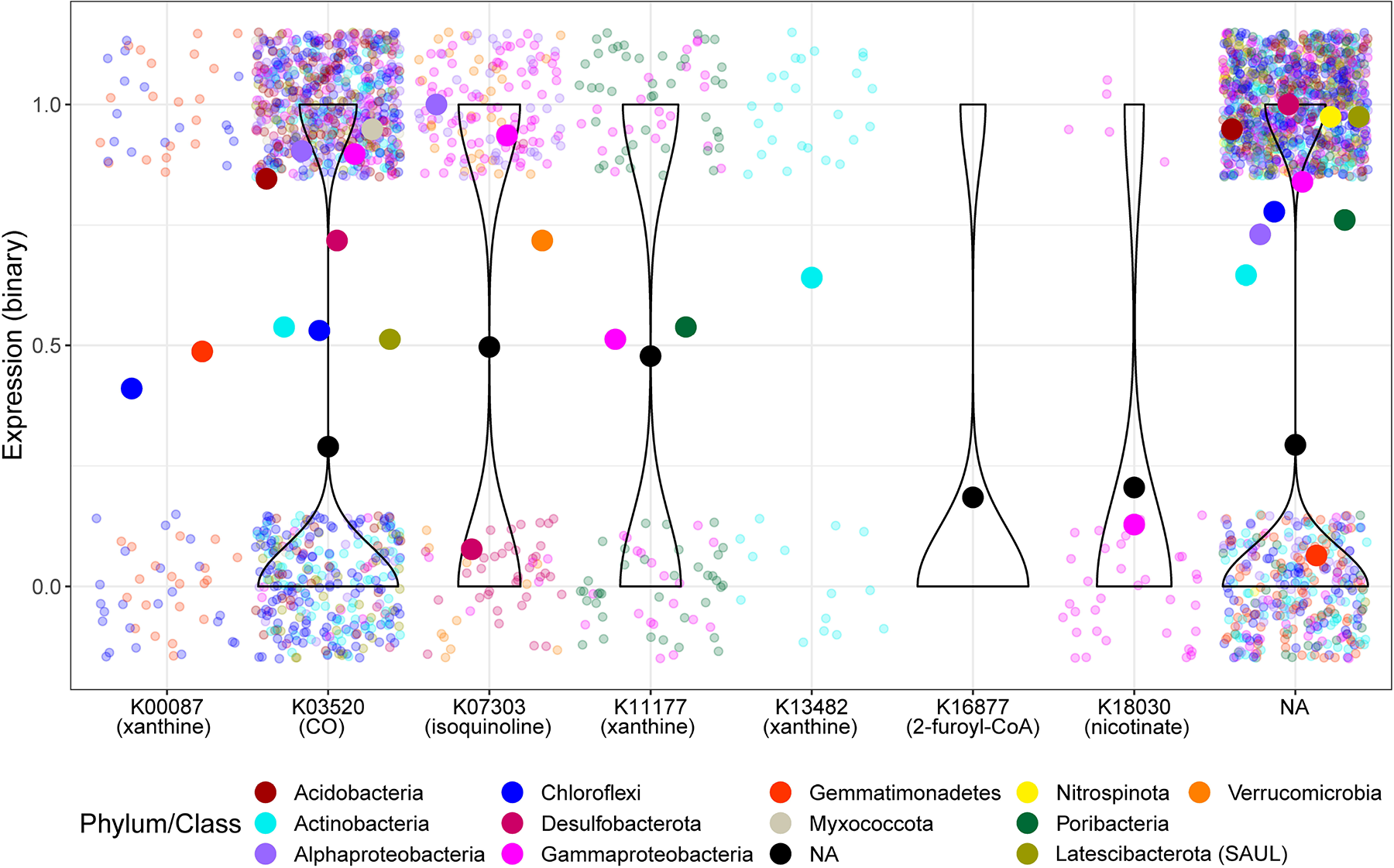
Expression of COG1529 and the variety of its hypothesized target molecules across phyla/classes of sponge symbionts in *P. ficiformis*. The analyses are based on cumulative binary (1 – expressed, 0 – not expressed) expression of transcripts with the same function and taxonomy (transcripts with the same KEGG annotations and MAG affiliations are merged) related to the different transcripts annotated as COG1529 across 39 samples of *P. ficiformis*. Taxonomy of transcripts is assigned if the transcript was linked to the gene of the assembled MAG. The potential target molecule is written in brackets. Larger dots represent proportion of expression across samples (average values from the binary data) for a certain taxonomy group (Phylum/Class). Violin plots (B and C) represent distribution of transcripts with no assigned taxonomy (NA).

It has been suggested that CODH supplies energy for enhanced anaplerotic reactions by pyruvate carboxylase (PYC) in planktonic marine Alphaproteobacteria [3]. Anaplerotic carbon assimilation may contribute differently to the actual biomass accumulation ranging 0.5–1.2% of the total carbon of cells [8] and 10-15% of protein [94] in different Alphaproteobacteria strains. Due to a high abundance of genes related to anaplerotic reactions (86%) within the fifty *P. ficiformis*-derived MAGs, we next determined the taxa that consistently expressed genes associated with anaplerotic carbon assimilation across multiple *P. ficiformis* specimens. Consistent expression by the same taxon across different specimens implies an enhanced anaplerotic flow, which can result in actual carbon assimilation due to relatively high carbon influx to the TCA cycle, while sporadic expression is more likely to be related to metabolic flexibility with periodical replenishment of TCA intermediates [4]. We observed prevalent expression (≥ 90% of all samples, n=39) of transcripts showing high similarity to anaplerotic proteins (PYC, malic enzyme [MEZ], and phosphoenolpyruvate carboxykinase [PCKA]) belonging to 3 Acidobacteria (order *Vicinamibacterales*), one Alphaproteobacteria (order UBA2966), and one Chloroflexi (order UBA3495). Thus, anaplerotic carbon assimilation in *P. ficiformis* might occur in Acidobacteria, Alphaproteobacteria and Chloroflexi. When genes were mapped against metatranscriptome reads, only the MAG-specific affiliation of *pckA* from Acidobacteria (MAG Acido_2) was confirmed (Figure 7). We further observed correlations between expression levels of *coxL* and PCKA transcripts, linked to Acido_2 MAGs, across twelve different *P. ficiformis* samples (Figure S3). In contrast to PYC, phosphoenolpyruvate carboxylase (PPC) and MEZ, PCKA utilizes CO_2_ rather than bicarbonate. We hypothesize that in Acido_2 within *P. ficiformis* and, possibly, in closely related *Vicinamibacterales* symbionts of *I. ramosa* (Figure 1, Figure S2, Clade Acd1), inorganic carbon assimilation may occur by CoxL supplying CO_2_ to the anaplerotic reaction catalyzed by PCKA.

**Figure 7.**
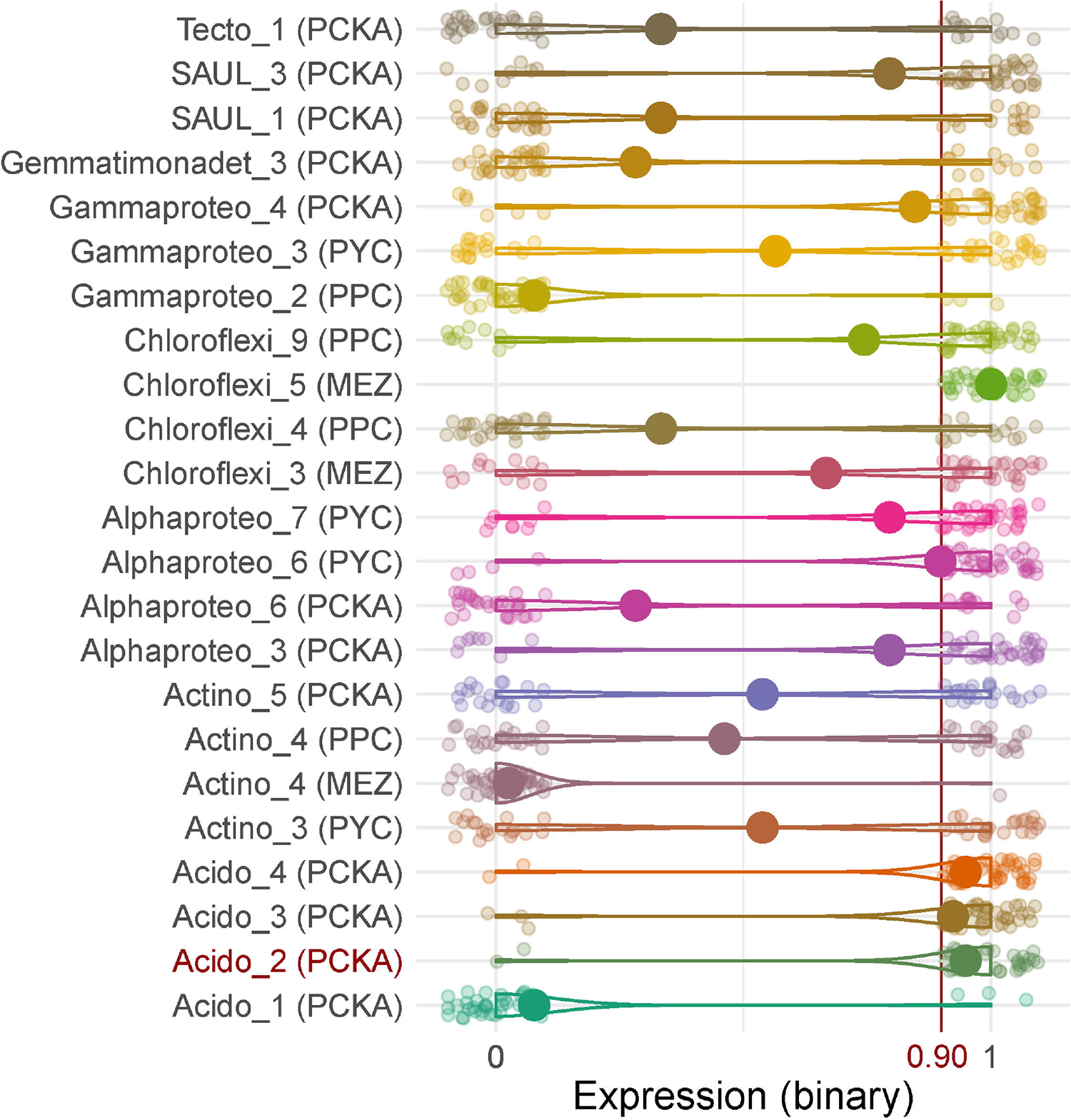
Anaplerotic carbon assimilation in *P. ficiformis* based on gene expression data (N=39). Cumulative binary expressions of transcripts related to anaplerotic carbon assimilation with identical functional (in bracket) and species (linked to a certain MAG) affiliations. Larger dots represent proportion of expression across samples. Genes that showed prevalent expression (>90% of samples) and their prevalence were also confirmed by direct mapping against the metatranscriptome reads (>90% of samples) are marked in red. MEZ, malic enzyme; PCKA, phosphoenolpyruvate carboxykinase; PPC, phosphoenolpyruvate carboxylase; PYC, pyruvate carboxylase.

Genomic potential for CBB was here found in symbionts of the Cyanobacteria and Proteobacterota (Gammaproteobacteria) phyla and was previously reported for Tectomicrobia [61]. Here we investigated the expression of the large subunit of RuBisCO (*rbcL*) in *P. ficiformis.* As expected, all samples that harbored Cyanobacteria (pink phenotype [31], n=12) showed expression of *rbcL*. In addition, we observed expression of a gammaproteobacterial *rbcL* in 30 (out of 39) samples (Figure 5C). This suggests that the Italian population of *P. ficiformis* (used for the transcriptomics data) is associated with a specific gammaproteobacterial symbiont with CBB activity, providing capability for dark fixation, while the Israeli population of *P. ficiformis* (used for obtaining MAGs) appears to lack this symbiont. A biogeographic effect on the microbial composition of *P. ficiformis* was reported before [32, 38].

Microbial carbon fixation can also occur through the 3-HP/4-HB cycle, which was suggested to be energetically fueled by ammonia oxidation in sponge-associated archaea [36]. Thaumarchaeota MAGs from *P. ficiformis* harbored *amoABC* genes (Figure 1, Table S4) and expressed *amoC* (Figure S4), as well as acetyl-CoA/propionyl-CoA carboxylase, the key gene of the 3-HP/4-HB cycle (Figure 5C). These findings confirm the involvement of Thaumarchaeota in dark carbon fixation in this sponge species. Orthologues of *amoABC/pmoABC* were also found in Desulfobacterota from *P. ficiformis* and are attributed to methane oxidation as explained below. The *pmoA* subunit of this MAG was expressed in 37 out of the 39 *P. ficiformis* samples suggesting a wide distribution for methane oxidation in *P. ficiformis* (Figure S4A.)

### Carbon fixation measurements in sponges

Physiology experiments, using ^14^C labelled bicarbonate can test the ability of autotrophic carbon assimilation that is light dependent (photosynthetic activity) or that occurs in darkness (dark primary production). Two sponge species were used in the ^14^C labelled bicarbonate fixation experiments: (1) *P. ficiformis,* harboring *Ca.* S. feldmannii, and (2) *T. swinhoei,* with *Ca.* S. spongiarum. The latter sponge is also known to harbor a dense population of filamentous Tectomicrobia that have genomic potential to fix carbon *via* CBB, utilizing multiple inorganic energy sources (Table S4, Figure 1).

Light-mediated inorganic carbon fixation was detected in both species in the cortex (external layer) of the sponge, where Cyanobacteria reside. While on average *ca*. 81-97% of total (*i.e.* light+dark) carbon fixation occurred in light conditions in *P. ficiformis* (Figure 8B,D), the overall contribution of light fixation in *T. swinhoei* ranged between 46 and 78% (Figure 8A, C). Transfer of labelled photosynthates to internal sponge layers was only observed for *T. swinhoei* (Figure 8A). In contrast, the labeled photosynthates produced by *Ca*. S. feldmannii remained within the cortex of *P. ficiformis* (Figure 8B, D). The lack of contribution of fixed organic carbon from *Ca*. S. feldmannii to internal layers of the sponge host supports the previous hypothesis that the symbiotic role of this photosymbiont may not be directly related to its photosynthetic properties, and rather to protection from solar radiation via synthesis of pigments [31]. Accordingly, presence of a photosymbiont does not directly imply transfer of organic carbon to its host, and alternative benefits need to be investigated. Diverse trends in carbon contribution to the host were shown also for different sponge species harboring *Ca.* S. spongiarum, and it was suggested that such variability may relate to symbiont phylotypes (clades within *Ca.* S. spongiarum) [19, 26].

**Figure 8.**
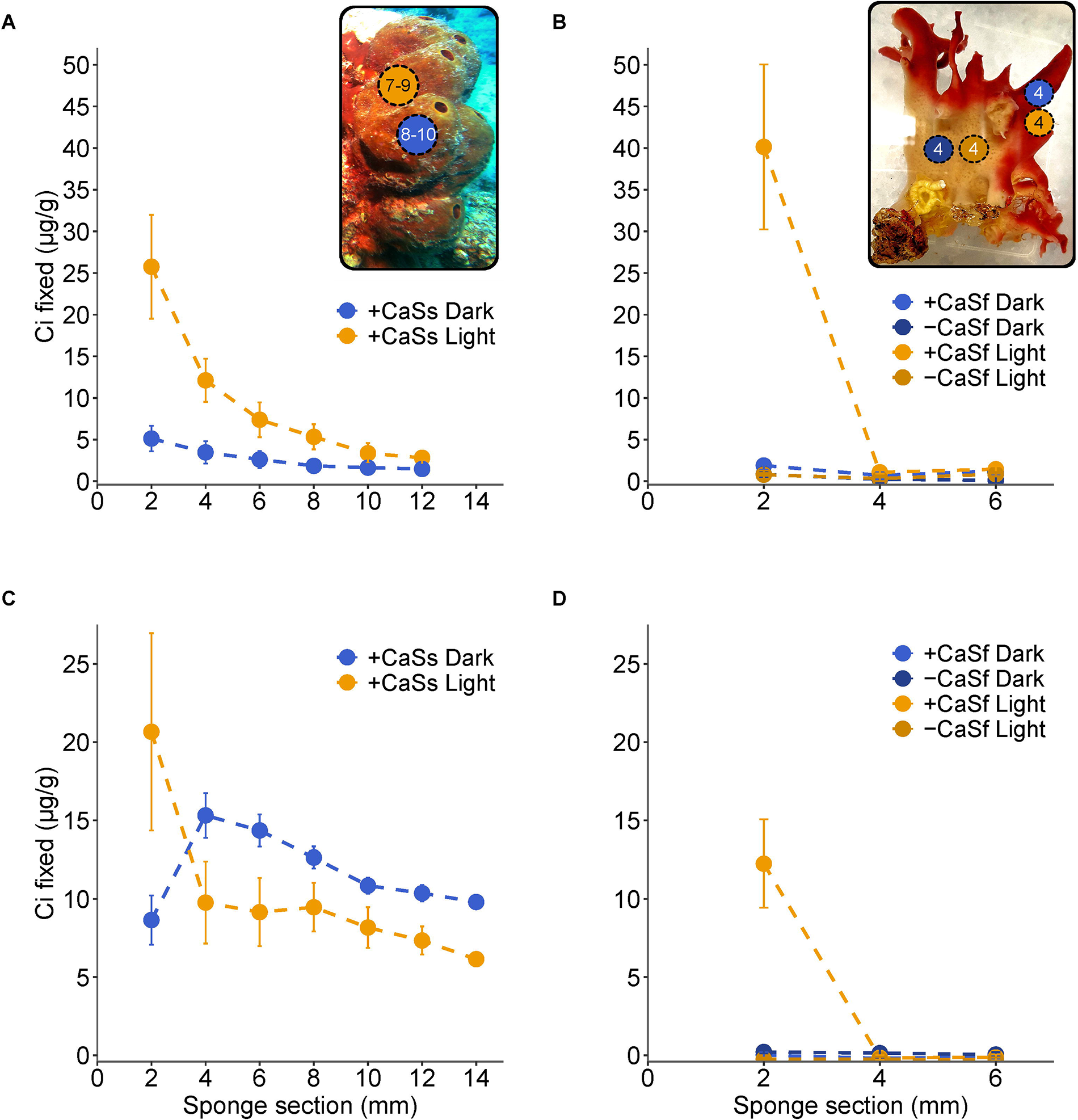
Light and dark carbon fixation in *T. swinhoei* (left insert) and *P. ficiformis* (right insert). In inserts, circles schematically represent cores with numbers of replications. Amounts (µg/g) of fixed labeled Ci across parallel sections of the sponge, including the most external (outer 2 mm, harboring cyanobacterial symbionts) and internal (cyanobacteria-free) sections. (A and C) Two specimens of *T. swinhoei* were used for each experiment, one was incubated in light and one in dark for two hours. At the end of the incubation, cylinders were cut out of the sponge and divided into 6-7 sections to establish labeled carbon fixation. Mean ± SD (n= 9 (A), and n= 7 (C) for light and n=10 (A), and n= 8 (C) dark conditions). CaSs, *Ca*. S. spongiarum. (B and D) Cores derived from light (cyanobacteria harboring) and dark (cyanobacteria-free) exposed parts of one specimen of *P. ficiformis* for each experiment were incubated for two hours with H^14^CO_3_^-^, cores were then cut and divided into 3 sections to establish labeled carbon fixation. Mean ± SD (n=4 for light and n=4 for dark conditions). CaSf, *Ca*. S. feldmannii.

Chemosynthetic or dark carbon fixation in *T. swinhoei* represented 16.6-29.5% of total fixation. Besides symbiotic Thaumarchaeota [38] and Nitrospirota [95]*, T. swinhoei* is also known to harbor abundant filamentous Tectomicrobia (‘Entotheonella’) This symbiont may therefore be the main organism responsible for the observed dark-fixation in this sponge species, by means of CBB cycle, fueled by a wide range of inorganic energy sources.

In contrast to *T. swinhoei*, dark fixation provided a relatively low contribution to total fixation (0.1-4.5%) in *P. ficiformis*. This, despite that *P. ficiformis* harbors Nitrospirota [31] and Thaumarchaeota [31, 38] symbionts, phyla that we show here as capable of dark carbon fixation via the rTCA and 3-HP/4-HB cycles, respectively. Both cycles are energetically fueled by different stages of nitrification, with ammonia and nitrite oxidation processes driven by Thaumarchaeota and Nitrospirota, respectively. However, ammonia oxidation rates were shown to be ten times lower than nitrite oxidation rates in the Mediterranean sponges *Dysidea avara* and *Chondrosia reniformis* [96], suggesting a larger influence of the rTCA compared to the 3-HP/4-HB cycle towards dark carbon fixation. If a similar trend is relevant also to *P. ficiformis*, then Thaumarchaeota may contribute little fixed carbon, resulting in the very low dark fixation observed in the ^14^C-label experiments. Nitrospirota were reported to have low relative abundance and activity in *P. ficiformis* [31], which can also explain their low impact to the overall carbon fixation in the holobiont. Finally, non-photosynthetic CBB fixation by Gammaproteobacteria, shown in this study for Italian *P. ficiformis* specimens based on detection of *rbcL* transcripts, may not be relevant in Israeli *P. ficiformis* specimens, where homologues of the same *rbcL* gene were not detected in metagenomes or MAGs (Figure 1, Figure S6). Taken together with the ^14^C-label experiments conducted on Israeli *P. ficiformis* specimens, chemoautotrophic pathways have only a minor influence on the overall carbon fixation compared to the photoautotrophic activity of *Ca.* S. feldmannii.

A decreasing pattern in H^14^CO_3_^-^ concentration in the medium in which the *P. ficiformis* cores were incubated was observed for both the Cyanobacteria-harboring cores (where H^14^CO_3_^-^ was fixed by *Ca*. S. feldmannii) (Figure S5A, B), and the white cores without Cyanobacteria (Figure S5A, C). Killed samples (formalin controls) did not show decreasing patterns of H^14^CO_3_^-^ in the incubation medium (Figure S5), implying biologically active uptake in living cores in the dark as well as the light. Given the minimal dark fixation observed, we suggest that the uptake of H^14^CO_3_^-^ by the sponge in the dark resulted from assimilation of bicarbonate via anaplerotic reactions followed by immediate respiration of most of the assimilated carbon to CO_2_.

If the dark-fixed H^14^CO_3_^-^ was indeed immediately respired back to CO_2_, we should only have detected the decrease in labelled H^14^CO_3_^-^ in the seawater if it had remained trapped inside the sponge tissue. We thus conducted an additional experiment with white (Cyanobacteria-free) *P. ficiformis* cores incubated with H^14^CO_3_^-^ in the dark, and once the decrease of labelled H^14^CO_3_^-^ in the medium was detected, we crushed the sponge tissue. This resulted in an increase of label in the medium indicating release of the labeled CO_2_ from the sponge core back to the medium (Figure S5C). This supports a fast turnover of dark-fixed CO_2_ in *P. ficiformis,* which might be related to anaplerotic carbon assimilation. Further, based on our results, anaplerotic carbon assimilation in *P. ficiformis* likely results in energy production rather than in biomass accumulation.

The anaplerotic rates in the laboratory conditions may be lower than in the natural environment, due to differences in accessibility to metabolically important compounds [97] such as electron donors (*e.g.*, pelagic CO). In fact, physiological experiments performed on planktonic Gammaproteobacteria showed increased rates of anaplerotic Ci assimilation when the appropriate energy source (*e.g.*, thiosulfate) and anaplerotic carbon acceptor (*e.g.*, pyruvate) were added [2]. We therefore cannot exclude that anaplerotic carbon assimilation in laboratory conditions might be different from the natural conditions.

## Conclusions

We have shown that CO oxidation is ubiquitous in sponge symbionts, likely representing the main inorganic energy source for lithoheterotrophs. Different variations of CODH and *amoABC*/*pmoABC* found across symbiotic lineages have evolved towards oxidation of diverse inorganic (*e.g.*, CO and ammonia) and organic (*e.g.*, xanthine and methane) compounds, that may be dissolved in seawater, that is continuously pumped through the sponge water channels, or produced within the sponge holobiont. Anaerobic forms of CODH and the WL pathway, previously suggested as being part of the Ci-fixing metabolic repertoire of some sponge symbionts, were found to be absent from the sponge microbiome. Most sponge symbionts were found to be lithoheterotrophs or organoheterotrophs with the exception of taxonomically restricted groups of autotrophs that implement the 3-HP/4-HB, CBB, and rTCA pathways. We provide the first experimental evidence for dark fixation in sponges, particularly in *T. swinhoei*. We further suggest that dark fixation processes in *P. ficiformis* (and potentially other sponge species) may also involve anaplerotic carbon assimilation, which is likely carried out by Acidobacteria, and possibly also by Alphaproteobacteria and Chloroflexi. Finally, cyanobacterial *Parasynechococcus*-like symbionts were shown to be highly diverse in terms of their contribution to the overall holobiont carbon budget, with *Ca.* S. spongiarum sharing its photosynthates with the host, while *Ca.* S. feldmannii behaving as a ‘selfish’ guest.

## Acknowledgments

The authors thank the Inter-University Institute (IUI) in Eilat, Israel, for their technical support in SCUBA dives and laboratory availability for physiology experiments. LS wishes to warmly thank Prof. Sven Beer and Prof. Micha Ilan for support with the experiments on *T. swinhoei.* Dr. Eyal Rahav and Dr. Natasha Belkin are thanked for advice on radioactive measurements and calculations. Dr. Stefan Green, director of the DNA Services Facility at the University of Illinois at Chicago (UIC) is thanked for useful comments and suggestions on sequencing strategies. The authors also thank Igor Chebotar, high performance computing system (HPC) administrator of the faculty of natural sciences at University of Haifa for his technical support in software and hardware assistance.

## Data availability

MAGs from this study can be found under NCBI bioprojects ID PRJNA515489 (*P. ficiformis*), PRJNA255756 (*T. swinhoei*), PRJNA712987 (*A. aerophoba*) and PRJNA273429 (*I. variabilis*).

