## Supplementary File S2 and Supplementary Figures for "Rethinking symbiotic metabolism: trophic strategies in the microbiomes of different sponge species"

#### Supplementary methods

##### *Taxonomy affiliation of the assembled scaffolds*

Taxonomy affiliation of separate scaffolds derived from the assembly of the sample 277c [1] was done as previously described [2] and using MEGAN-LR algorithm [3] that was done as follows. The whole scaffold sequences were searched against the NCBI-NR database (downloaded at 25/09/2018) with DIAMOND using the long reads mode (blastx --long-reads). The resulted data file was "meganized" using standalone daa-meganizer (--minScore 50 --longReads --maxExpected 0.001 --minPercentIdentity 45 --topPercent 10 --minSupportPercent 0.05 --minSupport 0 --lcaAlgorithm weighted --lcaCoveragePercent 80) and the protein accession to taxonomy prot_acc2tax-Nov2018X1.abin file. The taxonomy binning of the scaffolds was than accessed using MEGAN v6.15.2.

##### *Binning of MAGs using VizBin, differential coverage and ESOM*

Preliminary genomic binning with VizBin (default parameter), [4] resulted in 25 MAGs (Table S2). Manual binning procedure of additional 25 MAGs involved three steps and was conducted as follows. First, we calculated differential coverages for scaffolds assembled from the 277c sample [1] based on mapping against three *Petrosia ficiformis* samples [277c

(SAMN11333419), 287ce (SAMN12828169), 288c (SAMN11333441)] and plotted this data on 3D graph (File S1). We identified potential MAGs as scaffold subsets that clustering together ("clouds"). We draw an additional graph for every "cloud" or a group of "clouds" with similar coverage, taxonomy or GC% content patterns. These "clouds" - initial bins were extracted and cleaned from potential contamination using pentanucleotide frequencies-based clustering with VizBin.

***Annotation and clustering of CODH subunits***

In the wide genomic analysis Mo-CODH complex was annotated based on the presence of four subunits (CoxM COG1319, CoxL COG1529, CoxS COG2080, and CoxG COG3427) [5]. Custom HMM profiles were used to confirm presence and absence of Mo-CODH and Ni-CODH, respectively (Table S2). Parallel annotation using KEGG was done to clarify functional diversity of COG1529. 'mmseqs easy-cluster' (MMseqs2 version 12.113e3) [6] with default parameters was used to group enzymes annotated as COG1529 into clusters. Resulted clusters were visualized using Gephi [7].

***RuBisCO large subunit (*rbcL*) sequences in sponge microbiome***

Homology sequences for the putative gammaproteobacterial transcript from Italian *P. ficiformis* annotated as RuBisCO (TRINITY_DN30221_c0_g1_i1) were searched using BLASTX 2.11.0+ [8] against the NCBI-nr and using CD-Search [9] against the CDSEARCH/cdd [10] databases. *rbcL* domain of the transcript was extracted and used for the further analyses including BLASTN 2.11.0+ [11] against NCBI-nt. The extracted domain sequence was also aligned with the *rbcL* sequences obtained in 3 gammaproteobacterial MAGs (assembled from the sponge *I. ramosa*), *Ca*. S. feldmannii genomes and one derived from the assembled data of the 277c *P. ficiformis* specimen using BLASTN 2.11.0+.

We mapped reads from 44 sponge metagenomes derived from 6 sponge species (Table S5) against the custom database included 3 *rbcL* sequences obtained in 3 MAGs (*I. ramosa*) and transcript from Italian *P. ficiformis* using bbmap tool v 37.62 [12] from the BBtools package (https://jgi.doe.gov/data-and-tools/bbtools/). Counts per million reads was calculated for metagenomes with ≥ 10 read pairs mapped to the gammaproteobacterial *rbcL*.

All but 4 metagenomes were quality trimmed (quality threshold 20) with the reformat tool from the BBtools. 3300003175 (IMG accession number) metagenome was quality trimmed as described in [2]. The other three (277c, 287ce, 288c) were quality trimmed as described above. Transcript of cyanobacterial RuBisCO large subunit (*rbcL*) was confirmed using BLASTN 2.2.30+ [11] by searching of *rbcL* gene of *Ca*. S. feldmannii against assembled 160,550 transcripts [13] (identity=99.65%, *E*-value=0, score=2577).

***Taxonomical analysis of the phylum Tectomicrobia (`Entotheonellaeota`)***

16S rRNA gene sequences of Tectomicrobia (termed `Entotheonellaeota` based on 16S SILVA annotation) were obtained from previous studies and included 1 sequence from *P. ficiformis* [13], 125 sequences derived from various sponge species (including *T. swinhoei*), seawater and soil [14, 15], and 3 sequences assembled from the metagenomes 277c, 287ce, and 288c [1, 16]. These near-full-length 16S rRNA gene sequences were obtained from the metagenomic data and annotated as described in [17]. The resulted 129 sequences and 2 additional genes of Nitrospina (AM110965 and L35504) were aligned using the SILVA SINA aligner [18]. Additional closely related 45 `Entotheonellaeota`16S rRNA gene sequences from the SILVA database were incorporated to the alignment. Final alignment contained 176 nucleotide sequences and total of 236 positions. The final Maximum Likelihood tree (with Nitrospina as an outgroup) based on the Kimura 2-parameter substitution model [19] and discrete Gamma distribution that was used to model evolutionary rate differences among sites (5 categories (+G, parameter = 0.7137)) was constructed with MEGA7 [20]. Phylogenetic robustness was inferred from 1,000 bootstrap replications.

***Sponge sampling for the carbon fixation measurements***

Sponge samples were collected by SCUBA diving: three *P. ficiformis* specimens were collected at 22-27.5 m depths at Achziv nature marine reserve, Mediterranean Sea, Israel, on May 13th and June 9th, 2020; four specimens of *T. swinhoei* were collected at 25-32 m depths in Eilat, Gulf of Aqaba, Red Sea, Israel in 2000. One *P. ficiformis* specimen was held in the aquarium for 47 days prior to the carbon fixation experiment (Figure 8). Aquaria conditions were: 12/12 hours light regime (15 μE $m^{-2}$ $s^{-1}$), at 22 C. Sponge samples were collected in compliance with the permits from the Israel Nature and National Park Protection Authority.

***Core preparation and calculations of fixed carbon***

Cores of sponge tissue were cut perpendicularly to the surface. The surface area of each core was ~0.79 $cm^2$ and its depth ranged of 0.8 – 1.6 cm. After incubation with labeled bicarbonate, each core was cut into 2mm sections that are parallel to the sponge surface. These sections were weighed and transferred to the separate scintillation vials.

The amount of fixed carbon (X, in µg) was calculated as follows:

$$X = \frac{(\mathrm{DPM} - \mathrm{kDPM}) \cdot \mathrm{Vf} \cdot \mathrm{W} \cdot \ \mathrm{F} \ \cdot \ 1000}{\mathrm{tDPM}}$$

Where, DPM is the average DPM of three replicates, kDPM are the counts measured for the negative control (sponge core that was killed by exposure to formalin prior to incubation); Vf is the volume factor (here 6, as 100 µl were counted out of 600 µl); W is the estimated mass of carbon in seawater and equals 25 mg/L; F is the fractionation factor (uneven uptake of $^{14}C$ and

$^{12}C$) and equals 1.05; tDPM reflects the specific activity, average DPM counts (N=4) for $NaH^{14}CO_3$ in 100 µl of incubation medium, 1000 is the conversion from mg to µg.

### Supplementary text

#### *Large RuBiSCO subunit (*rbcL*) in Cyanobacteria and Gammaproteobacteria*

The *rbcL* domain of the assembled gammaproteobacterial transcript (TRINITY_DN30221_c0_g1_i1) was linked to the *rbcL* gene from genomes of *Sulfuricaulis limicola* and *Sulfurifustis variabilis* (identity=82-83%, cover=99-100%, *E*-value=0), which are sulfur oxidizers that were isolated from a lake in Japan [21, 22]. The gammaproteobacterial MAGs assembled from *P. ficiformis* lacked RuBiSCO subunits. To investigate whether the gammaproteobacterial RuBiSCO was not binned and assembled as a part of MAGs or was not present in the Israeli specimens of *P. ficiformis*, we first looked for all assembled *rbcL* sequences in 277c. Then we compared the sequence of the *rbcL* domain that was expressed in Italian *P. ficiformis* specimens with those assembled in Israeli *P. ficiformis* and in *I. ramosa*. Surprisingly, we found that the *rbcL*-domain sequence from Italian *P. ficiformis* showed higher similarity to the sequences derived from the three Gammaproteobacteria from *I. ramosa* (identity=75-76%, cover=100 %, *E*-value=0) than to three *rbcL* genes assembled from the metagenomes of the Israeli *P. ficiformis* specimens (data not shown). Thus, proteobacterial chemoautotrophs utilizing CBB cycle might exist in Israeli *P. ficiformis*, but probably be taxonomically different from the Italian *P. ficiformis* and from *I. ramosa*. To support this finding, we mapped 47 metagenomes representing six sponge species against the combined dataset of gammaproteobacterial *rbcL* sequences from Italian *P. ficiformis* and *I. ramosa* (Figure S6). The RuBiSCO type derived from the Italian transcriptome was absent from the three Israeli *P. ficiformis* specimens (277c, 287ce, and 288c).

Italian *P. ficiformis* may harbour Gammaproteobacterial symbionts taxonomically closely related to those assembled from *I. ramosa*. The distribution of these symbionts might be affected by biogeography [23]. However, not all *I. ramosa* samples harboured this type of RuBISCO with 25% of the samples did not pass the threshold of ≥20 mapped reads. We conclude that the distribution of this symbiont across sponge populations might be mosaic.

***Detailed discussion about taxonomical and morphological diversity of Tectomicrobia in sponges***

Based on 16S rRNA analyses, Tectomicrobia consists of three distant subclades`, with the most studied one being the filamentous so-called ‘Entotheonella’ from *T. swinhoei*, with four MAGs available [24–26]. Tectomicrobia are also present in *P. ficiformis* [13], however, they are taxonomically distant from the Tectomicrobia (supported by 98% bootstrap), which are found in *T. swinhoei*. Large filaments, which is the typical morphology of ‘Entotheonella’ symbionts, have never been reported in the microscopic studies of *P. ficiformis* [23, 27, 28], thus we speculate that Tectomicrobia associated with this sponge might be single celled.

***Methane monooxygenases (pmoABC) in Desulfobacterota***

Ammonia and methane monooxygenases (*amoABC/pmoABC*) are known for their similarity with each other and even the ability to oxidize both substrates [29, 30]. We affiliated *amoABC/pmoABC* of Desulfobacterota to methane oxidation due to its high similarity to the previously annotated methane monooxygenases from unclassified Deltaproteobacteria (nr NCBI databases, WP_066887011, identity=85%, *E*-value=3e-166) [31] and from *Streptomyces thermoautotrophicus* (RefSeq Select proteins databases, MBI3798512, identity=64%, *E*-value=1e-109).

***Nitrite oxidoreductase and nitrate reductase in Alphaproteobacteria, Gammaproteobacteria and Nitrospirota***

A previous genomic study on the *I. ramosa* microbiome proposed the presence of genomic ability for nitrate reduction via the nitrate reductase complex NarGHI in Alphaproteobacteria, Gammaproteobacteria and Nitrospirota sponge symbionts [32]. We here confirmed the presence of the first stage of denitrification (*narGHI* complex) in thirteen MAGs belonging to Alphaproteobacteria and Gammaproteobacteria phyla derived from *A. aerophoba*, *P. ficifirmis*, *I. ramosa*, and *Spongia officinalis*. However, in Nitrospirota MAGs, instead of nitrate reductase, we found nitrite oxidation genomic potential through the nitrite oxidoreductase *nxrAB* complex. *nxrA* and *narG* both have the KEGG annotation K00370, and *nxrB* and *narH* both belong to K00371. Because these genes belong to the same orthologous groups, additional genomic identifiers are required to distinguish between these functions in these MAGs. Nitrospirota MAGs lacked the third *narI* subunit (K00374), which is required for the NarGHI complex formation. By using specialized HMM profiles we confirmed the presence of nitrite oxidoreductase (*nxrAB*) in Nitrospirota (Table S1, Table S4). Altogether, the first stage of denitrification is performed by Alphaproteobacteria and Gammaproteobacteria MAGs. Nitraspirota have the capability for the last nitrification stage.

***CO sources for sponge symbionts***

Sponge-associated CO-oxidizing bacteria might obtain CO from two main sources. One being the non-biologically, photoproduced CO derived from the ambient seawater [33, 34] thanks to the water-pumping activity of the sponge [35]. Another being through biological hemoprotein degradation via heme oxygenase (HO) activity, as previously shown for symbionts associated with humans [36] and legumes [5, 37]. Sponge symbiotic bacteria may produce CO by metabolism of heme- and non-heme-related compounds from seawater DOM accumulated by host [36, 38].

Hemoproteins can also be biosynthesized by microbial symbionts inside the sponge. Heme synthesis and export was here found to be widespread among sponge symbionts with a total of 139 MAGs (including representatives of Acidobacteria, Actinobacteria, Chloroflexi, Gammaproteobacteria, Gemmatimonadetes and Nitrospirota) that contained both heme synthesis (protoheme synthase *hemBCDHYE* and/or siroheme synthesis *hemBCD*, *cysG*) and heme exporter genes (*ccmABC*) (Table S4). Other symbionts can then oxidize this synthesized and exported heme *via hemO* and *hguZ* [36], which were here annotated in 35 MAGs (including Acidobacteria, Alphaproteobacteria, Gammaproteobacteria and Tectomicrobia) (Table S4), resulting in CO production.

CO concentrations are known to affect external electron transport (EET) [39]. For example, in legumes, host-derived hemoprotein oxidation is responsible for the degradation of plant-produced leghaemoglobin, resulting in increased CO levels [5, 37], which, in turn, inhibit nitrogen-fixing rates of carboxydovore bacteria (bacteroids) associated to legumes [5]. Thus, by means of regulation of leghaemoglobin transfer to the bacteroids, the plant can control bacteroid metabolism [5]. We here suggest that fluctuations in CO concentrations inside sponges could also regulate global metabolism of the sponge microbiome. CO concentration in sponges will depend on various factors, including uptake, synthesis and degradation of this molecule. The sponge may alter CO concentrations through changes in water-pumping activity that concentrates DOC and supplies oceanic CO to its microbiome. The symbionts may affect CO concentrations inside the sponge through hemoprotein degradation (which is both part of the DOC as well as could be synthesized by the symbionts) and through CO oxidation.

In conclusion, CO levels, in a similar way to the legume symbiotic system, could affect EET in diverse members of the sponge microbiome and be related to the regulation of the symbiotic metabolism by both host and microbiome. This hypothesis is so far only based on gene content of the sponge microbiome community and requires to be tested by complementary methods.

15. Haygood DJFMG. Identification of the antifungal peptide-containing symbiont of the

marine sponge *Theonella swinhoei* as a novel Gammaproteobacterium , '*Candidatus* Entotheonella palauensis'. *Mar Biol* 2000; **136**: 969–977.

23. Burgsdorf I, Erwin PM, Lopez-Legentil S, Cerrano C, Haber M, Frenk S, et al. Biogeography rather than association with cyanobacteria structures symbiotic microbial

communities in the marine sponge *Petrosia ficiformis*. *Front Microbiol* 2014; **5**: 529.

30. Lontoh S, DiSpirito AA, Krema CL, Whittaker MR, Hooper AB, Semrau JD. Differential inhibition in vivo of ammonia monooxygenase, soluble methane monooxygenase and membrane-associated methane monoxygenase by phenylacetylene. *Environ Microbiol*

2000; **2**: 485–494.

38. Rix L, Ribes M, Coma R, Jahn MT, de Goeij JM, van Oevelen D, et al. Heterotrophy in the earliest gut: a single-cell view of heterotrophic carbon and nitrogen assimilation in sponge-

microbe symbioses. *ISME J* 2020; 1751–7370.

39. Tanaka K, Yokoe S, Igarashi K, Takashino M, Ishikawa M, Hori K, et al. Extracellular electron transfer via outer membrane cytochromes in a methanotrophic bacterium *Methylococcus capsulatus* (Bath). *Front Microbiol* 2018; **9**: 1–7.

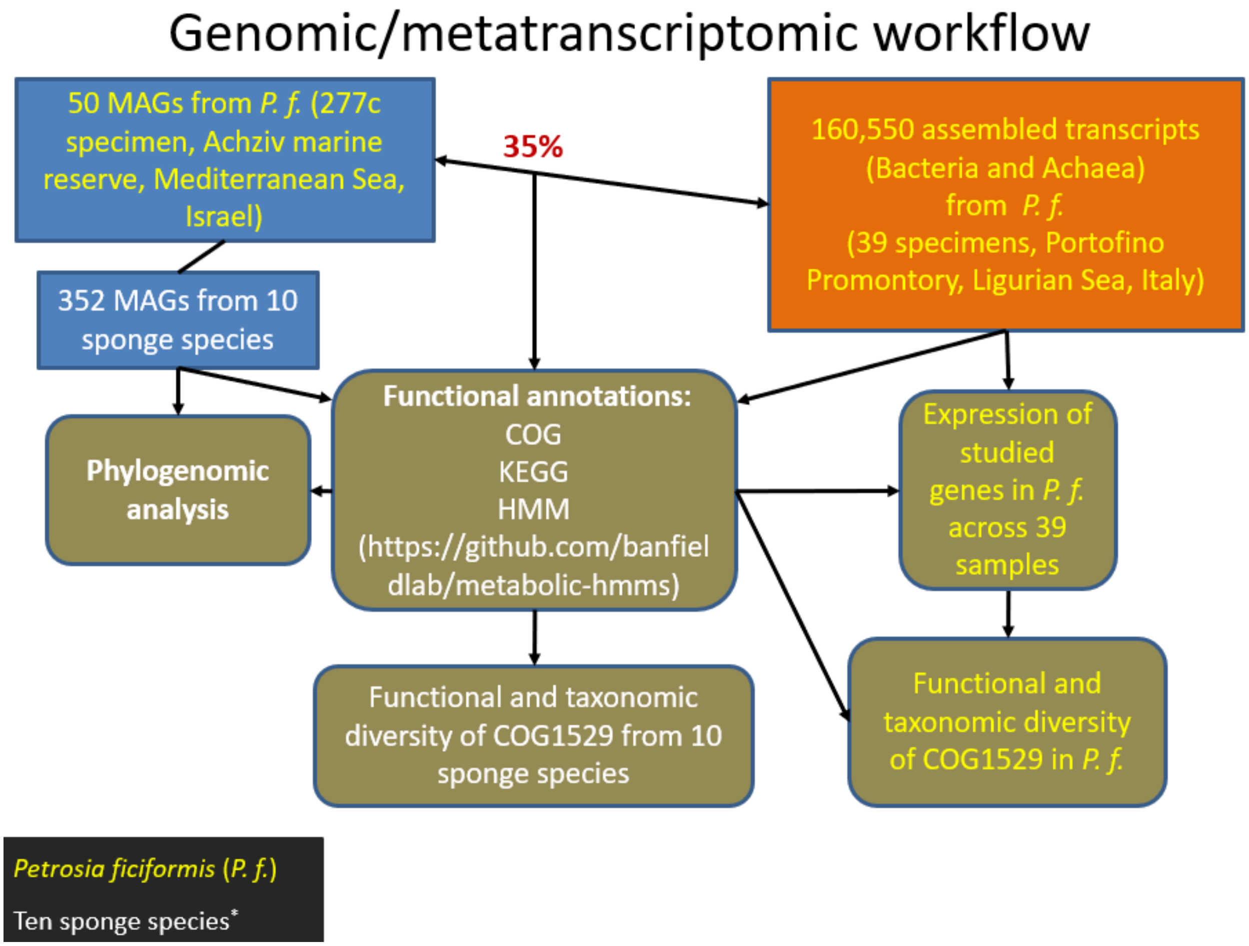


**Figure S1. Schematic representations of genomic and genome-centered metatranscriptomic analyses that were performed as a part of this study.** Additional ten sponge species including *Aplysina aerophoba, Carteriospongia foliascens, Dactylospongia elegans, Haliclona cymaeformis, Ircinia ramosa, Ircinia variabilis, Lamellodysidea herbacea, Petrosia ficiformis, Spongia officinalis, Theonella swinhoei.* P.f., *Petrosia ficiformis*.

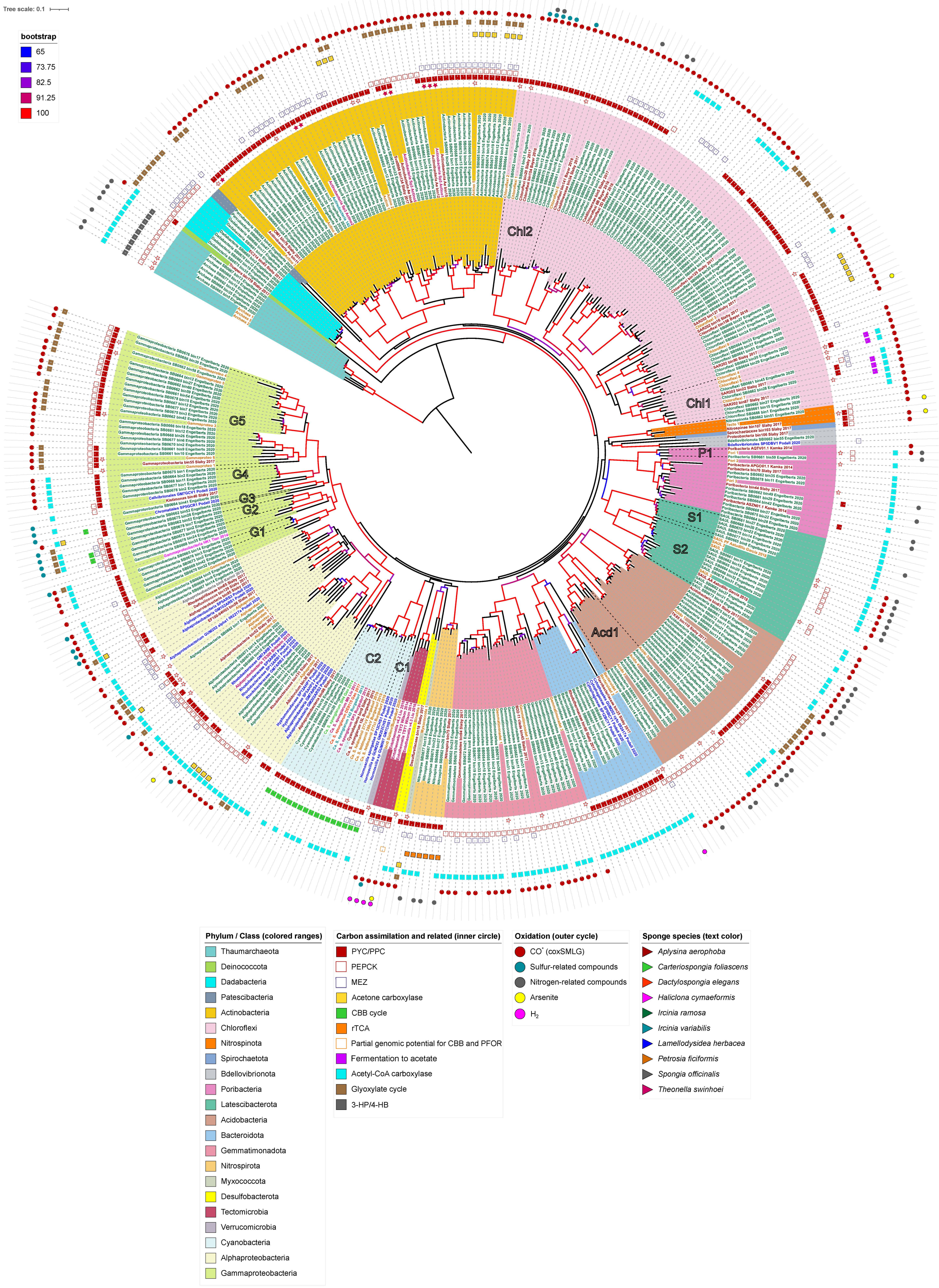
Tree scale: 0.1
bootstrap
65
73.75
82.5
91.25
100
Chl2
Chl1
P1
S1
S2
Acd1
C2
C1
G5
G4
G3
G2
G1
Phylum / Class (colored ranges)
Thaumarchaeota
Deinococcota
Dadabacteria
Patescibacteria
Actinobacteria
Chloroflexi
Nitrospinota
Spirochaetota
Bdellovibrionota
Poribacteria
Latescibacterota
Acidobacteria
Bacteroidota
Gemmatimonadota
Nitrospirota
Myxococcota
Desulfobacterota
Tectomicrobia
Verrucomicrobia
Cyanobacteria
Alphaproteobacteria
Gammaproteobacteria
Carbon assimilation and related (inner circle)
PYC/PPC
PEPCK
MEZ
Acetone carboxylase
CBB cycle
rTCA
Partial genomic potential for CBB and PFOR
Fermentation to acetate
Acetyl-CoA carboxylase
Glyoxylate cycle
3-HP/4-HB
Oxidation (outer cycle)
CO* (coxSMLG)
Sulfur-related compounds
Nitrogen-related compounds
Arsenite
H2
Sponge species (text color)
Aplysina aerophoba
Carteriospongia foliascens
Dactylospongia elegans
Haliclona cymaeformis
Ircinia ramosa
Ircinia variabilis
Lamellodysidea herbacea
Petrosia ficiformis
Spongia officinalis
Theonella swinhoei

**Figure S2. Phylogenomic tree (enhanced version of the Figure 1) showing the distribution and diversity of carbon assimilation and energy production pathways across microbial symbiont taxonomy and host-species**. The phylogenetic tree (N=399 MAGs) was constructed based on concatenated universal markers (PhyloPhlAn2). Labels marked with a hollow star are MAGs assembled in this study from the *P. ficiformis* specimen 277c. Labels marked with a colored star are eight MAGs assembled from the *A. aerophoba* specimen 15L, *T. swinhoei* specimen SP3 and *I. variabilis* specimen 142. The tree is rooted to the Archaea group. Acd1, class Vicinamibacteria, order *Vicinamibacterales*, family UBA8438. C1, order *Cyanobacteriales*, family *Desertifilaceae*. C2, order *Synechococcales*, family *Cyanobiaceae*. CHL1, class Dehalococcoidia, order UBA3495. CHL2, class Anaerolineae, order SBR1031. G1, order GCA-2729495. G2, order UBA10353, family LS-SOB. G3 (single MAG), order UBA4575. G4, order *Pseudomonadales*, *Pseudohongiellaceae* family. G5, order *Pseudomonadales*, HTCC2089 family. P1, class and order WGA-4E, unknown family. S1, unknown class. S2, UBA2968 class and order. *CO is not always a target molecule for the *coxSMLG* complex as it was shown here for *Poribacteria*.

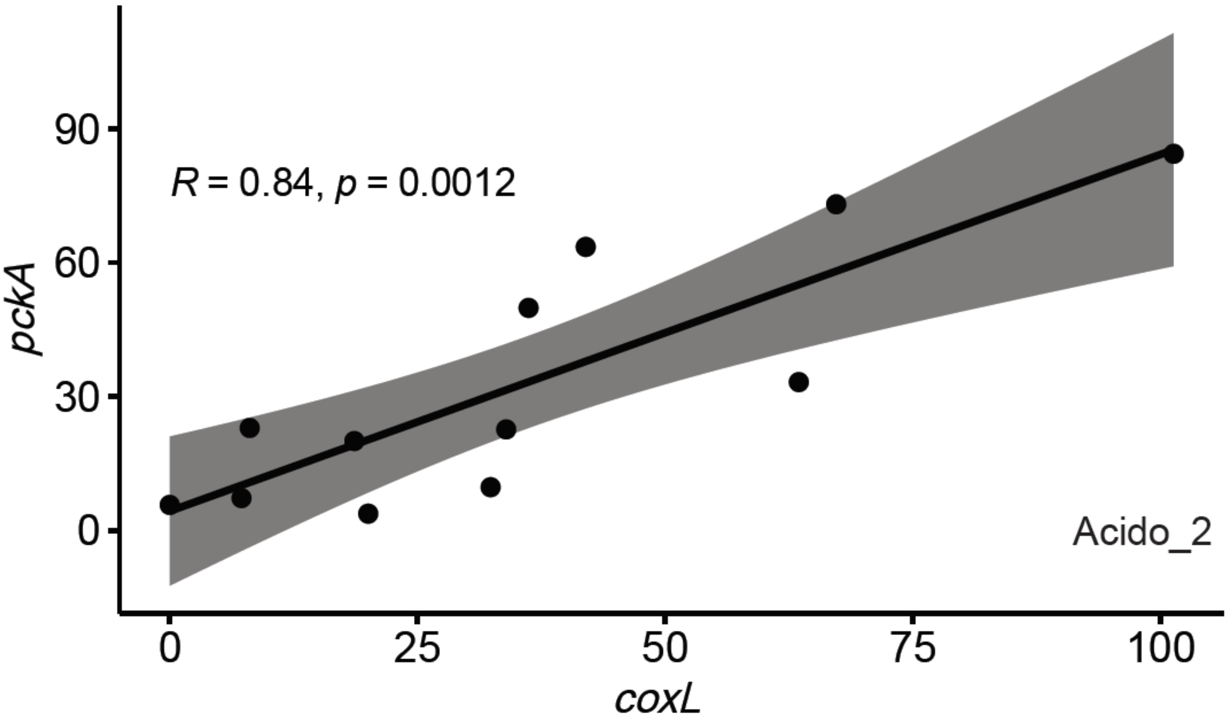


**Figure S3. Correlation of overall *coxL* and *pckA* expressions within the specific Acidobacteria symbiont Acido_2.** Spearman rank correlations of expression [number of aligned reads estimated by Salmon software (https://combine-lab.github.io/salmon/)] of *coxL* and *pckA* transcripts specifically linked to the Acidobacteria MAGs Acido_2 across twelve different *Petrosia ficiformis* samples. Separate transcripts with the same annotation (*coxL* or *pckA*) and belonging to the same MAG were merged prior to the analysis.

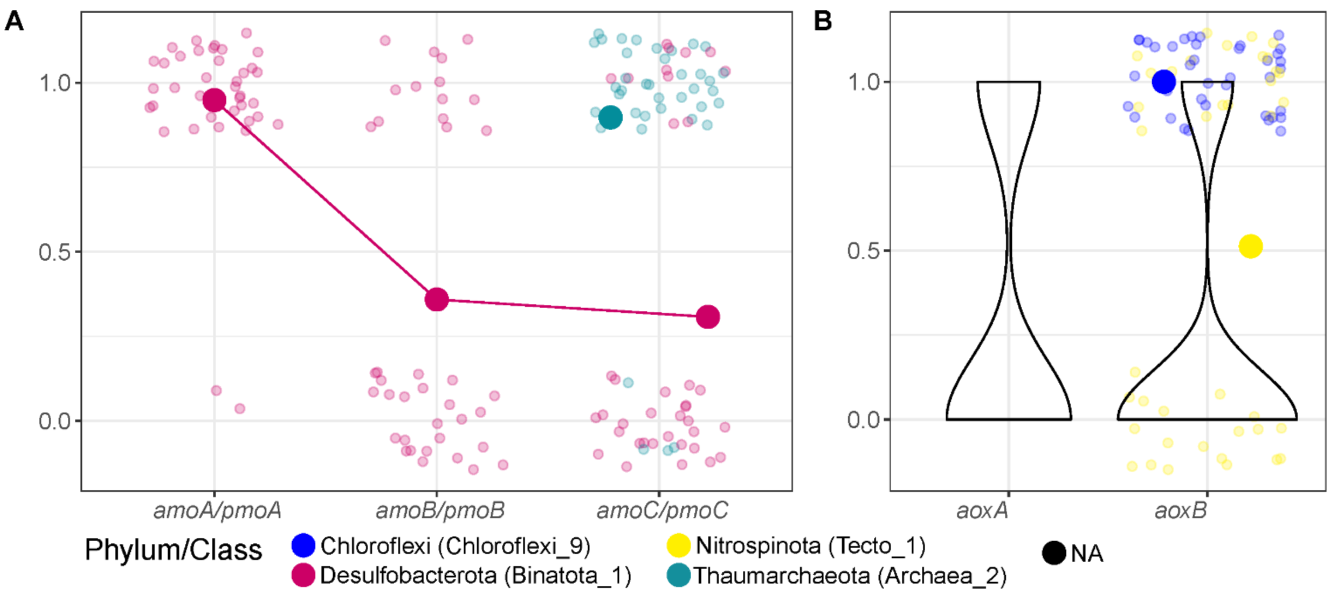


**Figure S4**. **Expression of ammonia/methane (*amoABC/pmoABC*) and arsenite (*aoxAB*) oxidation-related functions in the different phyla of *P. ficiformis* symbionts.** The analyses are based on cumulative binary (1 – expressed, 0 – not expressed) expression of transcripts (N=39 transcriptomes). Transcripts with the same function and MAG affiliations are merged. The three subunits of *amoABC/pmoABC* (subunits with the same taxonomy are connected by lines) (A) and *aoxB* subunit (B). Taxonomy of transcripts was assigned if the transcript was linked to the gene of the assembled MAG. Larger dots represent proportion of expression across samples for a certain taxonomy group (Phylum/Class). Transcripts with not assigned (NA) taxonomy (not linked to any assembled MAG) is given as a violin plot representing overall distribution of transcripts (B). The names of the MAGs with higher identity to the transcripts are presented in the brackets. *amoABC/pmoABC*, ammonia/methane monooxygenase; *aoxAB*, arsenite oxidase.

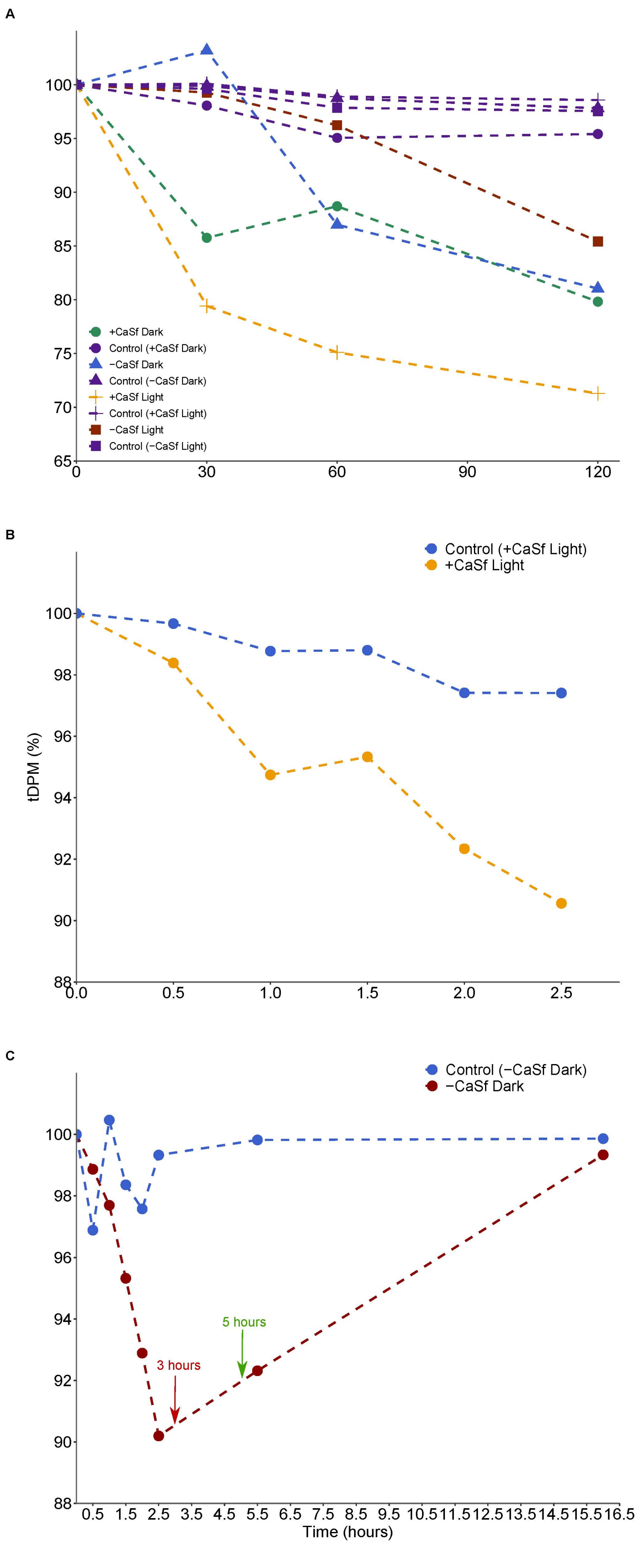
A
+CaSf Dark
Control (+CaSf Dark)
−CaSf Dark
Control (−CaSf Dark)
+CaSf Light
Control (+CaSf Light)
−CaSf Light
Control (−CaSf Light)
B
Control (+CaSf Light)
+CaSf Light
tDPM (%)
C
Control (−CaSf Dark)
−CaSf Dark
3 hours
5 hours
Time (hours)

**Figure S5**. **Labeled Ci uptake of *P. ficiformis* symbionts in light and dark conditions.** (A) Percentages of available labeled Ci per 0.1 ml of seawater in time (tDPM) in two beakers contained each four cores. (B) Measured tDPM in two beakers contained each two cores with *Ca*. S. feldmannii. (C) Measured tDPM in two beakers contained each four cores without *Ca*. S. feldmannii (white cortex). Red and green arrows represents the time points when the sponge tissue was disintegrated manually with a plastic homogenizer and N,N-Dimethylformamide was added, respectively. CaSf, *Ca*. S. feldmannii.

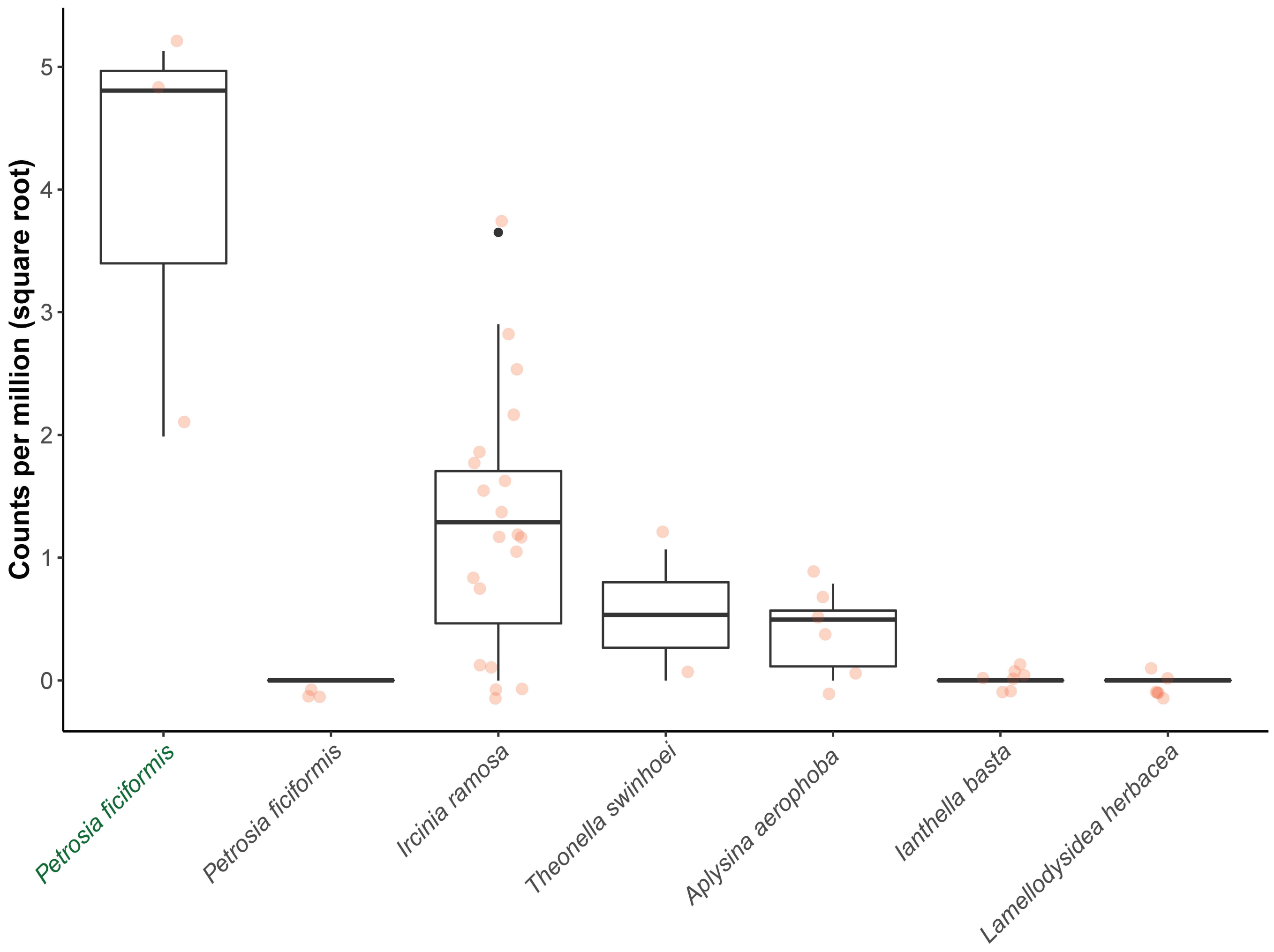


**Figure S6. Presence of large subunit of RuBiSCO (*rbcL*) derived from the Gammaproteobacteria symbionts in six sponge species**. Number of mapped reads (counts per million reads) against gammaproteobacterial (black font) and cyanobacterial (green font) *rbcL*. Gammaproteobacteria *rbcL* included three sequences from *Ircinia ramosa* (MAGs) and one from Italian population of *Petrosia ficifromis* (transcriptome). Cyanobacteria *rbcL* was taken from the genome of *Candidatus* Synecococcus feldmannii 277cV.
